# Optimizing Angiopep-2 Density on Polymeric Nanoparticles for Enhanced Blood-Brain Barrier Penetration and Glioblastoma Targeting: Insights from In Vitro and In Vivo Experiments

**DOI:** 10.1101/2024.11.05.622195

**Authors:** Weisen Zhang, Ahmed Refaat, Haoqin Li, Douer Zhu, Ziqiu Tong, Joseph A. Nicolazzo, Bo Peng, Hua Bai, Lars Esser, Nicolas H. Voelcker

## Abstract

The blood-brain barrier (BBB) poses a formidable challenge to efficient drug delivery into the brain. One promising approach involves leveraging receptor-mediated transcytosis facilitated by Angiopep-2 peptide (Ang-2)-conjugated nanoparticles. However, the precise impact of Ang-2 density on BBB penetration remains poorly understood. In this study, we developed a versatile polymeric nanoparticle system with tuneable Ang-2 surface density and systemically examined its influence on BBB penetration through various *in vitro* assays and an *in vivo* study. Our findings revealed a nuanced relationship between Ang-2 surface density and BBB penetration across the different experimental setups. In 2D cell culture, we observed a positive correlation between Ang-2 surface density and cellular association in hCMEC/D3 cells, characterized by a distinctive inflection point. Conversely, in the Transwell model, higher Ang-2 density correlated negatively with BBB penetration, whereas the BBB-GBM-on-a-chip showed the opposite trend. These disparities may arise from differences in avidity under static versus dynamic conditions, potentially modulating nanoparticle interactions due to fluidic forces. *In vivo* studies revealed that higher Ang-2 densities facilitated nanoparticle transport across the BBB, consistent with the findings of the BBB-GBM-on-a-chip model. Furthermore, loading doxorubicin into nanoparticles with optimal Ang-2 density resulted in controlled pH-responsive release and enhanced anticancer effect against U87 GBM cells in both 2D cell cultures and a 3D BBB-GBM-on-a-chip model. These results underscore the critical importance of optimizing Ang-2 surface density for efficient BBB penetration and emphasize the utility of dynamic models in preclinical *in vitro* assessment of novel nanoparticle formulations for targeted delivery to the brain.

## 1. Introduction

The brain homeostasis relies on a delicate balance of nutrients and neurotransmitters^1^. The supply and exchange of these chemicals to and from the brain is regulated by a unique property of the blood capillaries in the central nervous system (CNS): the blood-brain barrier (BBB)^2–4^. This selective barrier is comprised of a tight monolayer of brain endothelial cells maintaining proximity to glial cells and neurons, mediating communication between the CNS and the periphery^5, 6^. Potential treatments for CNS-related often do not progress to clinical trials and implementation due to the presence of the BBB, which often severely hinders the penetration of therapeutics into the brain. A promising strategy to overcome this delivery restriction is the use of the receptor-mediated transcytosis (RMT) pathway. This pathway involves a vesicular transcellular route through which macromolecules are transported across brain endothelial cells via ligand-receptor/protein binding, internalization, and releasing mechanisms^7^. Several target receptors on brain endothelial cells have been identified and explored for drug delivery, including the transferrin receptor^8^, insulin receptor^9^, glucose transporter^10^, and low-density lipoprotein receptor-related protein 1 (LRP1)^11^.

LRP1 is a critical transmembrane and ubiquitous signal protein that interacts with over 40 ligands, playing a pivotal role in regulating BBB integrity and functionality^12^. Notably, LRP1 is also highly expressed in glioblastoma (GBM)^13^, as it contributes to cancer invasion and migration^14^. This provides a rationale for designing nanoparticle systems targeting LRP1 to promote brain and GBM uptake. Angiopep-2 (Ang-2), a peptide derived from the Kunitz domain of human aprotinin^15^, exhibits a strong affinity to LRP1 and holds great potential in assisting nanoparticle delivery to the brain^16–19^. Among various types of nanoparticles, polymeric nanoparticles represent a promising class of nanocarriers due to their biocompatibility, tuneable structures, and ease of surface functionalization with targeting moieties^20^. Several studies have demonstrated the potential of using Ang-2 functionalized polymeric nanoparticles for CNS delivery^7, 21, 22^. However, the role of Ang-2 surface density in the ability of nanoparticles to cross the BBB remains ambiguous and some findings are contradictory. For instance, Tian *et al.* observed that polymersomes with the highest Ang-2 surface density (110 ligands per particle) showed significantly lower brain uptake in mice compared to the nanoparticles with lower Ang-2 surface density (1 and 22 ligands per particle)^23^. In contrast, Jiang *et al.* compared polystyrene nanoparticles coated with 10, 20, and 30 mol % of Ang-2, finding that higher Ang-2 coated nanoparticles led to higher BBB penetration^24^. Therefore, it is crucial to carefully evaluate and optimize the Ang-2 surface density on newly developed delivery systems to enhance their efficiency in achieving successful brain uptake.

Preclinical *in vitro* testing of bespoke nanoparticles for their permeability across the BBB is often performed using the conventional Transwell model, which lacks several key physiological features, including but not limited to, fluid flow. In this regard, microfluidic BBB-on-chip technology offers a more physiologically relevant platform for assessing permeability under hydrodynamic flow conditions. This key feature plays a critical role in endothelial cell proliferation and the dynamic interactions between nanoparticles and cells^25, 26^. The recent BBB-on-a-chip model, developed in our group, has been demonstrated to successfully visualize notable differences in the transportation of transferrin-coated nanoparticles, of different surface densities of transferrin, across the BBB ^27, 28^. Moreover, this BBB-on-a-chip platform can be integrated with a GBM cell line, such as U87 cells, to establish a more comprehensive *in vitro* BBB-brain cancer model, which was demonstrated to predict not only the permeabilities of nanomaterials across the BBB but also to monitor the transport of these particles in the brain cancer region^28^.

In this study, we developed polymeric nanoparticles with varying surface densities of Ang-2 to investigate their interaction with the BBB and identify the optimal surface density for brain targeting. Our research involved a comprehensive assessment of the fabricated nanoparticles using a combination of *in vitro* and *in vivo* assays, including cell association studies, studies of uptake pathway mechanisms, examination of BBB penetration in both static Transwell and dynamic microfluidic BBB models, as well as a brain uptake study in mice. The results provided valuable insights into how our nanoparticles behave in these different models, shedding light on the influence of Ang-2 surface density on BBB penetration. Subsequently, we explored the application of the optimized Ang-2 conjugated nanoparticles for targeted chemotherapeutic treatment of GBM. We introduced doxorubicin into the nanoparticle core via a Schiff-base reaction and evaluated its anticancer effects on GBM cells in both 2D cell cultures and 3D microfluidic BBB-GBM-on-a-chip models.

## 2. Results and discussion

### 2.1. Preparation and characterization of self-assembled polymeric nanoparticles (DAA100)

We synthesized a multifunctional self-assembled polymeric nanoparticle of controlled size using aqueous reversible addition-fragmentation chain transfer (RAFT)-mediated polymerization-induced self-assembly (PISA, **Figure 1A**). The composition of the prepared nanoparticle included surface azides for conjugation with the targeting ligand (Ang-2 DBCO), Cy5 fluorescent labels to facilitate *in vitro* and *in vivo* assays, and internal ketone groups for conjugation of an amine-containing drug (such as doxorubicin) via pH-responsive linkages. Additionally, a cross-linker was introduced to ensure nanoparticle stability after drug conjugation. Firstly, a macro-chain transfer agent (macro-CTA) was prepared by copolymerization of *N*,*N*-dimethylacrylamide (DMA), and the pre-synthesized Cy5-acrylamide (characterized by LC-MS with an m/z at 635.5 [M+H]^+^, **Figure S1**), and using an azido-functional chain transfer agent, 2-(dodecylthio-carbonothioylthio)-2-methylpropionic acid 3-azido-1-propanol ester. This yielded the macro-CTA P(DMA-*co*-Cy5) with a Mn of 5296 g/mol and a low dispersity (Đ) of 1.10 (**Figure S2**). The incorporation of Cy5-acrylamide was confirmed by SEC, showing a unimodal molecular weight distribution with correlated fluorescence emission (645 nm) at the same retention time as the polymer (**Figure S2B**).

**Figure 1.**
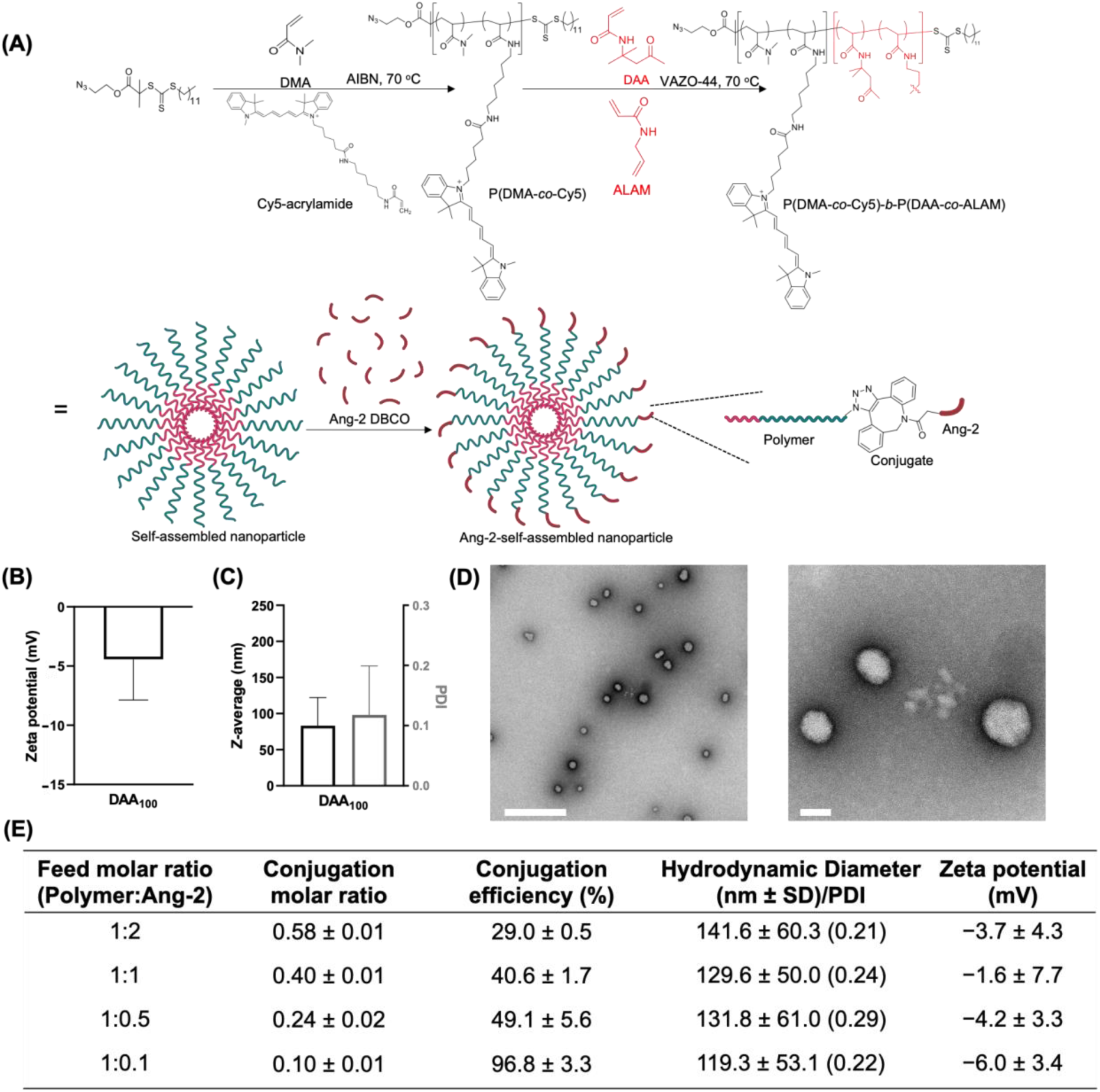
(A) Schematic diagram for the fabrication of azido-functionalized P(DMA-*co*-Cy5)-*b*-P(DAA-*co*-ALAM) polymeric nanoparticles (DAA_100_) and their conjugation to Ang-2 DBCO via a copper-free click reaction. (B) Zeta potential measurement. (C) Particle hydrodynamic diameter (ζ-average) alongside PDI, and (D) TEM images in low (left image, scale bar = 500 nm) and high (right image, scale bar = 50 nm) magnification. (E) Conjugation efficiency, hydrodynamic diameter, PDI, and zeta potential measurements of Ang-2-conjugated nanoparticles prepared at different molar feed ratios of azido-polymer:Ang-2 DBCO as 1:0.1, 1:0.5, 1:1, and 1:2. Conjugation efficiency was calculated based on a micro-BCA assay. Values in panels B, C and E are shown as mean ± SD (n = 3).

The self-assembled, core-crosslinked nanoparticles (DAA100) were prepared through aqueous dispersion polymerization of the macro-CTA with diacetone acrylamide (DAA) and the pre-synthesized cross-linker allyl acrylamide (ALAM) (**Figure S3**) using a molar feed ratio of DAA: ALAM: P(DMA-*co*-Cy5): VAZO-44 of 100: 1: 1: 0.2. This polymerization resulted in polymers production with an increased molecular weight (Mn = 82,593 g/mol) and a relatively high Đ of 1.80 due to core-cross-linking (**Figure S4**). ^1^H NMR spectra confirmed a successful chain extension with DAA (conversion >99%), as indicated by the characteristic methyl ketone proton signals of polymerized DAA at 2.20 ppm (f) (**Figure S5**). The incorporation of the cross-linker ALAM was further validated by evaluating the colloidal stability of the nanoparticles in a polar organic solvent (1,4-dioxane). Non-cross-linked self-assembled nanoparticles that were prepared with the same molar feed ratio of DAA were stable in water but disassembled in 1,4-dioxane, while the core cross-linked DAA100 retained a similar size and a polydispersity (PDI) in both water and 1,4-dioxane (**Figure S6**).

The morphology of the core cross-linked DAA100 nanoparticle was thoroughly characterized by dynamic light scattering (DLS) and transmission electron microscopy (TEM). DLS showed that DAA100 maintained a neutral charge and a hydrodynamic diameter of around 84.5 ± 38.9 nm with a polydispersity index (PDI) of 0.11 ± 0.08 (**Figure 1C**). TEM revealed that the nanoparticles had a spherical morphology with an average particle diameter of 64.6 ± 11.9 nm (n = 80 nanoparticles) (**Figure 1D**). The size discrepancy between DLS measurement and TEM imaging could be attributed to the different measurement environments as the TEM represents dried nanoparticles which might present some extent of shrinkage^29^. While not the primary emphasis of this study, it is important to note that the size of the nanoparticles can be readily adjusted by modifying the molar feed ratio of DAA^30^.

### 2.2. Polymeric nanoparticles (DAA100) with different surface densities of Ang-2

The DAA100 nanoparticles were modified with Ang-2 using copper-free click chemistry. A library of nanoparticles with a wide range of Ang-2 surface densities could be obtained by varying the molar feed ratio of the azido-group to the DBCO-modified Ang-2 from 1:0.1 to 1:2 (**Figure 1E**), leading to an conjugation efficiency from 10 to 58% as determined by a micro-bicinchoninic acid (micro-BCA) assay. As a complementary method, the amount of unreacted azido groups was determined using a fluorescence-based assay. Briefly, Ang-2-conjugated nanoparticles were solubilised in DMSO leading to dissolution of the polymers. This was followed by conjugating any unreacted azido groups with Cy3-DBCO which enabled back calculation to the conjugated molar ratios of Ang-2 in the different formulations. The latter mirrored the micro-BCA-derived data, further confirming the success of the conjugation method (**Figure S7**). Interestingly, the conjugation efficiency decreased significantly with increasing feed molar ratios of Ang-2. For instance, the 1:0.1 molar ratio (DAA100-Ang-2(1:0.1)) showed the highest conjugation efficiency (∼99%) while the 1:2 molar ratio resulted in the lowest conjugation efficiency. This might be caused by the steric hindrance effect between the conjugated Ang-2 on the surface of nanoparticles.

To estimate the number of Ang-2 molecules per nanoparticle or per unit surface area of nanoparticle (nm^2^), we first measured the number of nanoparticles per mg of DAA100 using a Nanosight Analyzer (4.5 × 10^12^ particles / mg) and estimated the surface area of nanoparticles based on their average hydrodynamic diameters. Then, we correlated the number of conjugated angiopep-2 molecules to the nanoparticles number or surface area as shown in **Figure S7**. The results indicated an increased number of Ang-2 molecules per nanoparticle from 319.5 ± 106.5 in DAA100-Ang-2(1:0.1) to 2067.1 ± 140.1 for DAA100-Ang-2(1:2). The number of Ang-2 molecules per nm^2^ of nanoparticle increased from 0.015 ± 0.003 to 0.091 ± 0.006 for DAA100-Ang-2(1:0.1) and DAA100-Ang-2(1:2), respectively. All Ang-2-conjugated DAA100 maintained an acceptable PDI of < 0.32 while their hydrodynamic diameters increased slightly from 119.3 ± 53.1 nm in the case of DAA100-Ang-2(1:0.1) to 141.6 ± 60.3 nm for DAA100-Ang-2(1:2) (**Figure 1E**). To further confirm the morphology of Ang-2 conjugated nanoparticles, we selected the nanoparticle with the highest Ang-2 density (DAA100-Ang-2(1:2)) for TEM analysis. The TEM of DAA100-Ang-2(1:2) displayed a spherical shape and an average particle diameter of 65.4 ± 12.8 nm (n = 80 nanoparticles), which was comparable to the control DAA100 (**Figure S8**). All samples maintained a close to neutral zeta potential range (+10 to −10 mV), potentially lowering the propensity for protein fouling^31, 32^.

### 2.3. Influence of Ang-2 surface density on nanoparticles association and uptake into human brain endothelial (hCMEC/D3) cells

The cytocompatibility of Ang-2 conjugated nanoparticles was first evaluated in hCMEC/D3 (**Figure S9**), and no significant cytotoxicity was observed across all the tested nanoparticle concentrations with the highest of 200 µg/mL. Subsequently, we employed confocal microscopy and flow cytometry to investigate the impact of Ang-2 surface densities on the association and uptake of the polymeric nanoparticles by endothelial cells. All the Ang-2-conjugated DAA100 showed higher Cy5 mean fluorescence signals in hCMEC/D3 cells, compared to the non-Ang-2 conjugated DAA100 (**Figure 2A**). This finding is consistent with previous research indicating the active role of Ang-2 in binding to LRP1 on brain endothelial cells, leading to increased uptake^33, 34^. The cellular uptake and internalization of Ang-2-conjugated nanoparticles in hCMEC/D3 cells were confirmed using z-stack scanning confocal microscopy. The images provided evidence that DAA100-Ang-2 (1:2) nanoparticles were located within the cells, suggesting a successful uptake and internalization of the nanoparticles (**Figure S10**).

**Figure 2.**
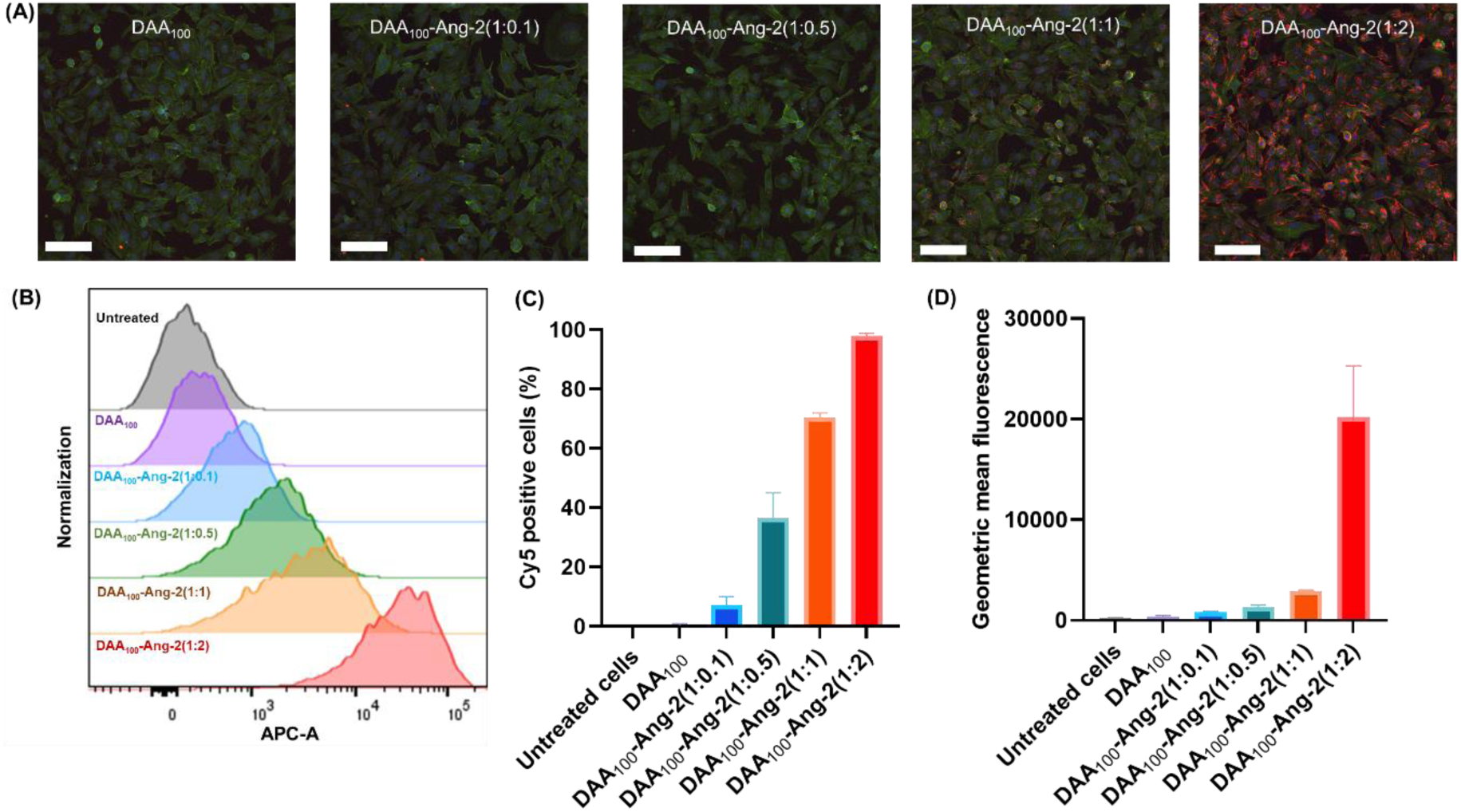
Cell association of different Ang-2-conjugated DAA_100_ to hCMEC/D3 cells. (A) Confocal fluorescence microscopy images of hCMEC/D3 cells after incubation with DAA_100_, DAA100-Ang-2 (1:0.1), DAA_100_-Ang-2(1:0.5), DAA_100_-Ang-2(1:1), and DAA100-Ang-2 (1:2). Nucleus is shown in blue, F-actin is shown in green, and Cy5 nanoparticles is presented in red. (scale bar = 50 µm). Flow cytometry analysis of the cellular association of different Ang-2-conjugated DAA_100_ to hCMEC/D3 cells (B) histogram, (C) percentage of Cy5 positive cells, and (D) the geometric mean fluorescence (GMF). Values are shown as mean ± SD (n = 3).

We performed a quantitative analysis of flow cytometry data to calculate the relative percentage of cells associated with nanoparticles. The percentage of Cy5-positive cells was compared with untreated cells, revealing a clear upward trend in cellular association as the Ang-2 density increased. This trend ranged from approximately 7.2 % for DAA100-Ang-2(1:0.1) to ∼97.8 % for DAA100-Ang-2(1:2) as shown in **Figure 2B** **and 2C**. The relative number of nanoparticles associated to cells was determined using geometric mean fluorescence (GMF), where the GMF of cells treated with DAA100-Ang-2(1:2) was ∼7 times higher than the second highest signal samples DAA100-Ang-2 (1:1) **(Figure 2D)**. In contrast, the GMF of cells treated with DAA100-Ang-2 (1:0.1) and DAA100-Ang-2(1:1) only increased from 812 to 2930. This result is consistent with the confocal microscopy images and suggests a positive correlation between the Ang-2 surface density and the nanoparticles’ cellular association to brain endothelial cells. A specific Ang-2 surface density threshold could exist for enhanced cellular association and uptake of nanoparticles^35^. This is probably attributed to varying strengths of binding interaction (avidity) to the LRP1 based on the Ang-2 surface density. For instance, Tian *et al.* reported a similar trend and combined this with a model based on super-selectivity theory ^36^. They found that the Ang-2 density on the polysomes posted a sigmoidal trend to the avidity of LRP1 with a selective peak at around 30 ligands per 50 nm-sized nanoparticle^37^. In addition, a dynamic increase in the expression level of LRP1 was found in response to the Ang-2 density by Liu *et al.* who demonstrated that Ang-2-coated PEG-PLGA nanoparticles induced the upregulation of LRP1 expression on epithelial cells to facilitate stronger absorption efficiency when compared to non-coated nanoparticles^38^.

### 2.4. Investigation of Ang-2-mediated uptake pathway in hCMEC/D3 cells

To further investigate how the Ang-2 surface density affects the mechanism of nanoparticle internalization, inhibitors of caveolae-mediated endocytosis (nystatin), clathrin-mediated endocytosis (chlorpromazine), and macropinocytosis (amiloride) were incubated with the hCMEC/D3 cells before their exposure to the nanoparticles. Firstly, none of the used inhibitors resulted in complete inhibition of particle uptake, indicating that Ang-2 conjugated nanoparticles likely employ more than one internalization pathway (**Figure 3**). Secondly, as the density of Ang-2 on nanoparticles increased, the cellular uptake was strongly reduced in chlorpromazine-pretreated cells. Given that clathrin-mediated endocytosis is the primary pathway responsible for receptor-mediated endocytosis^8^, this result reinforces the role of Ang-2 in facilitating nanoparticle receptor-mediated transcytosis for BBB transfer, consistent with other reports^34^. Moreover, the observed trend suggests an enhanced receptor-mediated endocytosis triggered by nanoparticles with higher Ang-2 surface densities.

**Figure 3.**
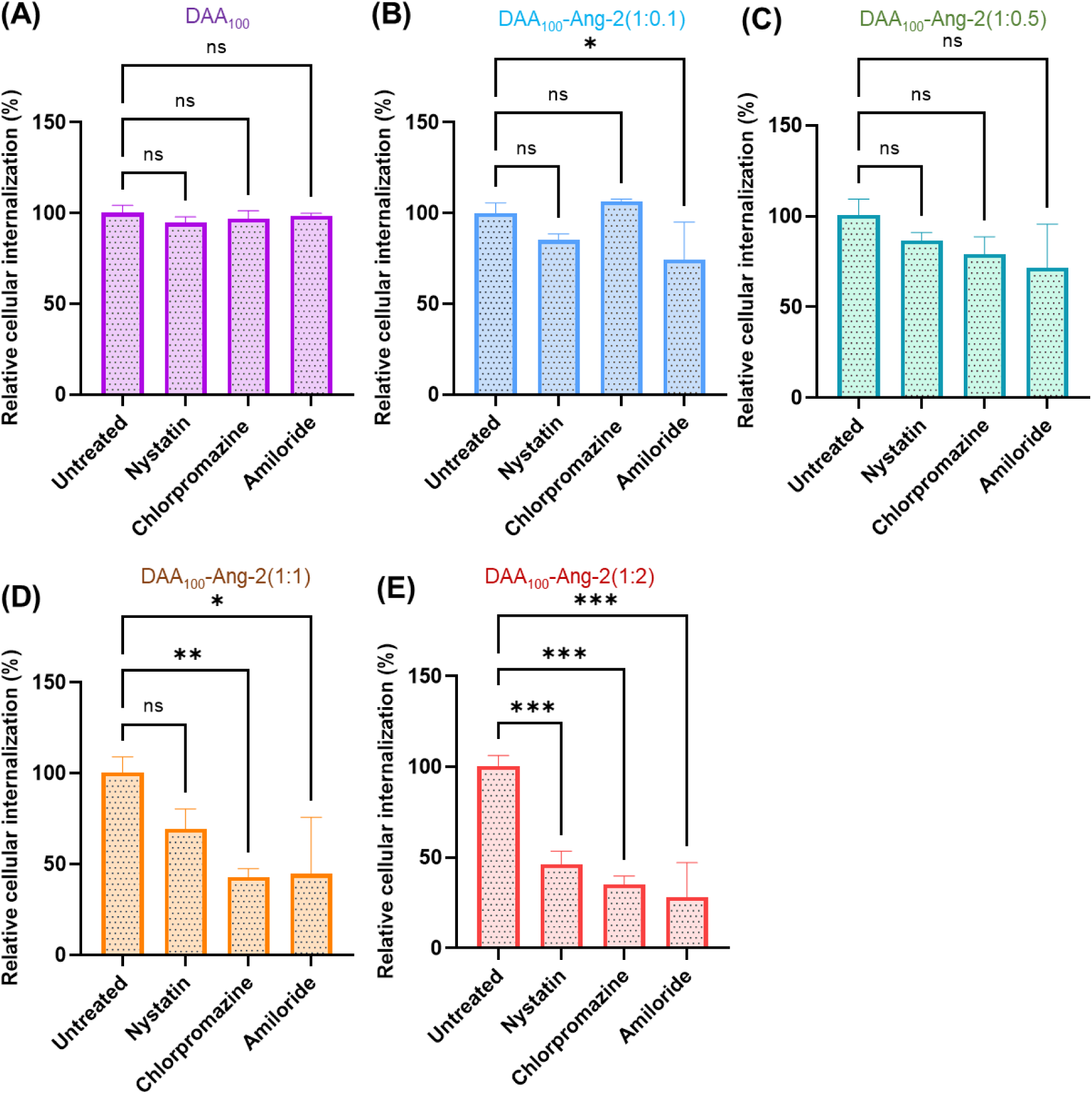
Influence of Ang-2 surface densities on nanoparticle internalization mechanism. Percentage of relative uptake of nanoparticles with different Ang-2 surface densities by hCMEC/D3 cells pre-treated with endocytosis inhibitors: nystatin (50 µg/mL), chlorpromazine (15 µg/mL), and amiloride (125 µg/mL). Percentage uptake of each nanoparticle type in the presence of inhibitors was determined relative to cells incubated with nanoparticles without inhibitors. Values are shown as mean ± SD (n = 3). One-way ANOVA with Tukey’s multiple comparisons test was used, * p < 0.05, ** p < 0.01, *** p < 0.001, **** p < 0.0001.

Interestingly, the macropinocytosis inhibitor (amiloride) and chlorpromazine shared a similar uptake reduction trend in the nanoparticles with higher Ang-2 surface density. Macropinocytosis is commonly regarded as non-selective cell “drinking” that results from actin-driven extension of plasma membrane ruffling in response to physiological cues (e.g. antigen, virus, soluble large molecule, etc.)^39, 40^. Our results suggest that higher Ang-2 surface density on the nanoparticles could increase non-selective “cell drinking”. This finding can be linked to the relationship between nanoparticle avidity and Ang-2 density, as aforementioned in section 2.3. Specifically, the avidity of LRP1 to the nanoparticles could significantly be enhanced by increasing the surface density of Ang-2. When LRP1 interacts with a higher density of Ang-2 on the surface of nanoparticles, the resultant strong adhesive nanoparticles could cause a membrane deformation, leading to “cell drinking”^23^.

Notably, only the nanoparticles with the highest Ang-2 densities DAA100-Ang-2(1:2) showed a significant reduction in cellular uptake when treated with nystatin (**Figure 3E**). Although caveolae-mediated endocytosis is not the main pathway responsible for receptor-mediated transcytosis, some reports reveal that caveolae-mediated endocytosis is involved in the brain endothelial cellular uptake of Ang-2 coated nanoparticles, as a parallel mechanism to clathrin-mediated endocytosis^41^. Our findings suggest the potential to determine a specific threshold of Ang-2 surface density on nanoparticles that can activate caveolae-mediated endocytosis. To trigger this pathway, the Ang-2 density of nanoparticles needs to exceed this defined range. This could be attributed to the fact that caveolae, not being primary structural elements for transcytosis, are more selective to specified binding ligands and rely on the integrity cytoskeleton^42^.

### 2.5. Study of nanoparticle permeability in a Transwell BBB model

The BBB transcytosis of nanoparticles with different Ang-2 densities was first assessed using a Transwell BBB model comprising a hCMEC/D3 cell monolayer. The hCMEC/D3 cells were seeded into the inserts, with 3 µm pore size, that separated the donor and receptor chambers and were cultured for 7 to 8 days until the trans-endothelial electrical resistance (TEER) values reached the characteristic range of 25-30 Ωcm^2^ (**Figure 4A** and **B**)^27, 43–46^. Afterward, Ang-2-conjugated nanoparticles were applied to the donor chamber, and the percentage of transported nanoparticles to the receptor chamber, relative to the initial amount, was quantified at 2, 4, 8, 24, and 48 h time points using fluorescence measurements.

**Figure 4.**
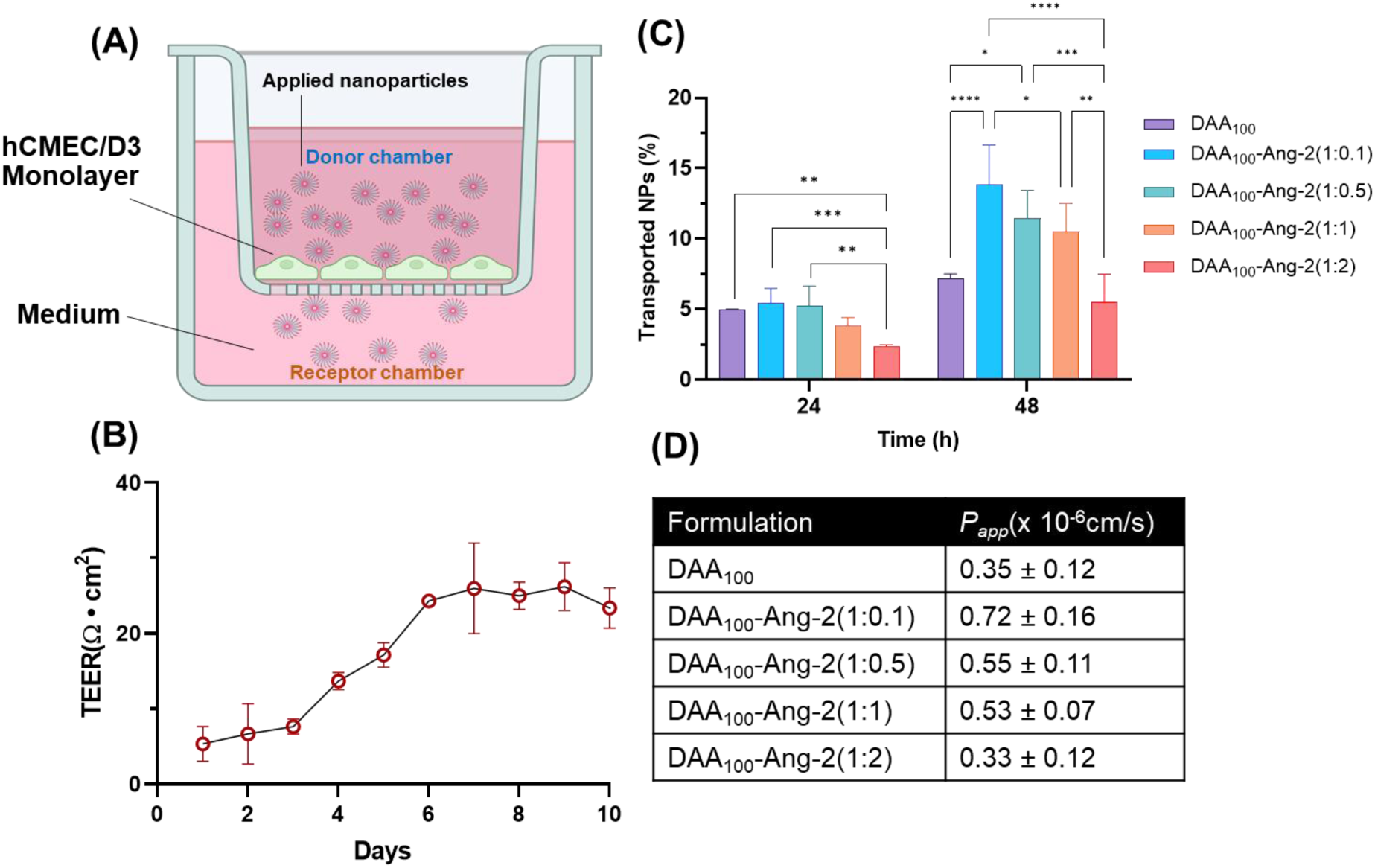
Nanoparticles permeability study in a Transwell BBB model. (A) Schematic representation of a BBB Transwell model that was formed by a monolayer of hCMEC/D3 cells over an insert with 3 µm pores. (B) TEER value measurements of hCMEC/D3 monolayer over 10 days. (C) Percentage of accumulated nanoparticles after 24 and 48 h in the receptor chamber of the BBB Transwell model after applying nanoparticles in the donor chamber. Two-way ANOVA test was used for data analysis with Tukey’s multiple comparisons test, * p < 0.05, ** p < 0.01, *** p < 0.001, **** p < 0.0001. (D) Apparent permeability coefficients (P_app_) of nanoparticles across the hCMEC/D3 monolayers in Transwell inserts. All data are shown as mean ± SD (n = 3).

In all treatment groups, the transported percentage of nanoparticles increased with time (**Figure S11**) and became most distinctive at 48 h (**Figure 4C**). Unlike the cellular association results, the transportation percentage showed an inverse correlation to the Ang-2 surface densities. DAA100-Ang-2(1:0.1) exhibited the highest transport (13.8 ± 2.7%), followed by DAA100-Ang-2(1:0.5) (11.4 ± 2.0%), DAA100-Ang-2(1:1) (10.5 ± 2.0%), DAA100 (7.17 ± 0.34%), and DAA100-Ang-2(1:2), (5.4 ± 1.9%). Remarkably, the penetration percentage of control nanoparticles (DAA100) was even higher than the nanoparticles with the highest Ang-2 content. Apparent permeability coefficient (*Papp*) values, representing the rate of transport across the barrier, were also calculated for all treatment groups (**Figure 4D**).

Ang-2-coated nanoparticles are expected to cross the BBB by binding to LRP1 on the endothelial cells and triggering receptor-mediated transcytosis. This receptor-mediated transcytosis comprises three steps: i) attachment of nanoparticles to LRP1 and subsequent endocytosis on the luminal side; ii) trafficking of nanoparticles within the cells, depending on the endocytic pathway involving movement of the carrier towards endolysosomal network; and iii) detachment of nanoparticles from the endosome and release to the other side of the membrane^23^. In principle, the binding interaction between nanoparticles and receptors should be strong enough to ensure sufficient duration for cargo uptake, yet not so overpowering that it prevents the release and transport of the cargo. As shown in sections 2.3 and 2.4, a higher Ang-2 density on nanoparticles promotes more attachment and uptake in the cells. However, the Transwell result indicated an inverse relationship between Ang-2 density and transport levels. This suggests an optimal threshold for surface ligand density, beyond which decreased BBB penetration would be observed. The underlying mechanism could be related to the avidity between nanoparticles and LRP1 or endosome. For instance, the avidity from the highest Ang-2 density could trigger a significantly strong attachment of nanoparticles but impede its trafficking and release, potentially leading to more degradation in lysosomes rather than transportation^47^. Tian *et al.* also demonstrated that Ang-2 density on the nanoparticles would trigger different LRP1 transcytosis pathways^23^. They found that high surface ligand density of cargo favors more lysosomal degradation over transcytosis. In contrast, a low or medium density of ligands helped nanoparticles towards a unique syndapin-2-related tubular deformation, promoting transportation rather than endolysosomal degradation and sorting. Such nonlinear dependence of ligand density and avidity on BBB crossing rate and related mechanism also correlates with results reported for the targeting of transferrin receptors^48–50^, glucose transporter-1^51^, and adhesion molecule-1^52^.

### 2.6. Evaluation of nanoparticle trafficking across a microfluidic BBB-GBM-on-a-chip model

While the Transwell assay provided some initial insights into nanoparticle penetration, it lacks the mechanical shear stress, which plays a vital role in endothelial cell proliferation^53–56^ and bio-nano interactions ^26^. To address this, a dynamic microfluidic *in vitro* model was used to investigate the effect of fluidic conditions on the BBB crossing ability of our Ang-2 polymeric nanoparticles and their subsequent uptake into GBM cells. We employed a BBB-GBM-on-a-chip model, which consisted of two main interconnected channels: the “blood” channel and the “brain” channel. The “blood” channel served as a simplified avatar of the BBB lined with hCMEC/D3 endothelial cells, while the brain channel contained U87 GBM cells to assess BBB nanoparticle penetration and targeting to GBM (**Figure 5A**). The two channels are interconnected via an array of microchannels (3 μm). This BBB-GBM model was established and validated in our previous reports^27, 28^. Besides the 3D cell culture capability and the fluid flow, our model allows for real-time monitoring of nanoparticles interaction with cells using high resolution fluorescence imaging^43, 57^. The fluorescence signal of associated Cy5 nanoparticles in both channels was measured to estimate the association of nanoparticles in endothelial cells and their BBB crossing potential (**Figure 5B** and **C**, representative images from chips perfused with DAA100-Ang-2(1:2) and DAA100-Ang-2(1:0.1)). Cellular associations of nanoparticles in the “blood” channel were compared to the signals from the control nanoparticles (DAA100) and shown as fold changes. Association of nanoparticles in the “blood” channel was positively correlating to their Ang-2 surface densities. Whereas relative to DAA100 control nanoparticles, DAA100-Ang-2(1:2) showed the highest cellular interaction followed by the ratios of 1:1, 1:0.5, and 1:0.1 (**Figure 5D**). A similar pattern was observed for the “brain” channel. Notably, DAA100-Ang-2 (1:2) showed a significantly higher signal (almost double) than DAA100-Ang-2(1:1) and ∼ 14 times higher than DAA100-Ang-2(1:0.1) (**Figure 5D**). The pattern of BBB transportation of the Ang-2 conjugated nanoparticles in the BBB-GBM-on-a-chip was significantly different from the Transwell result showing an almost reversed trend. This demonstrated the potential influence of a dynamic environment on BBB penetration of nanoparticles.

**Figure 5.**
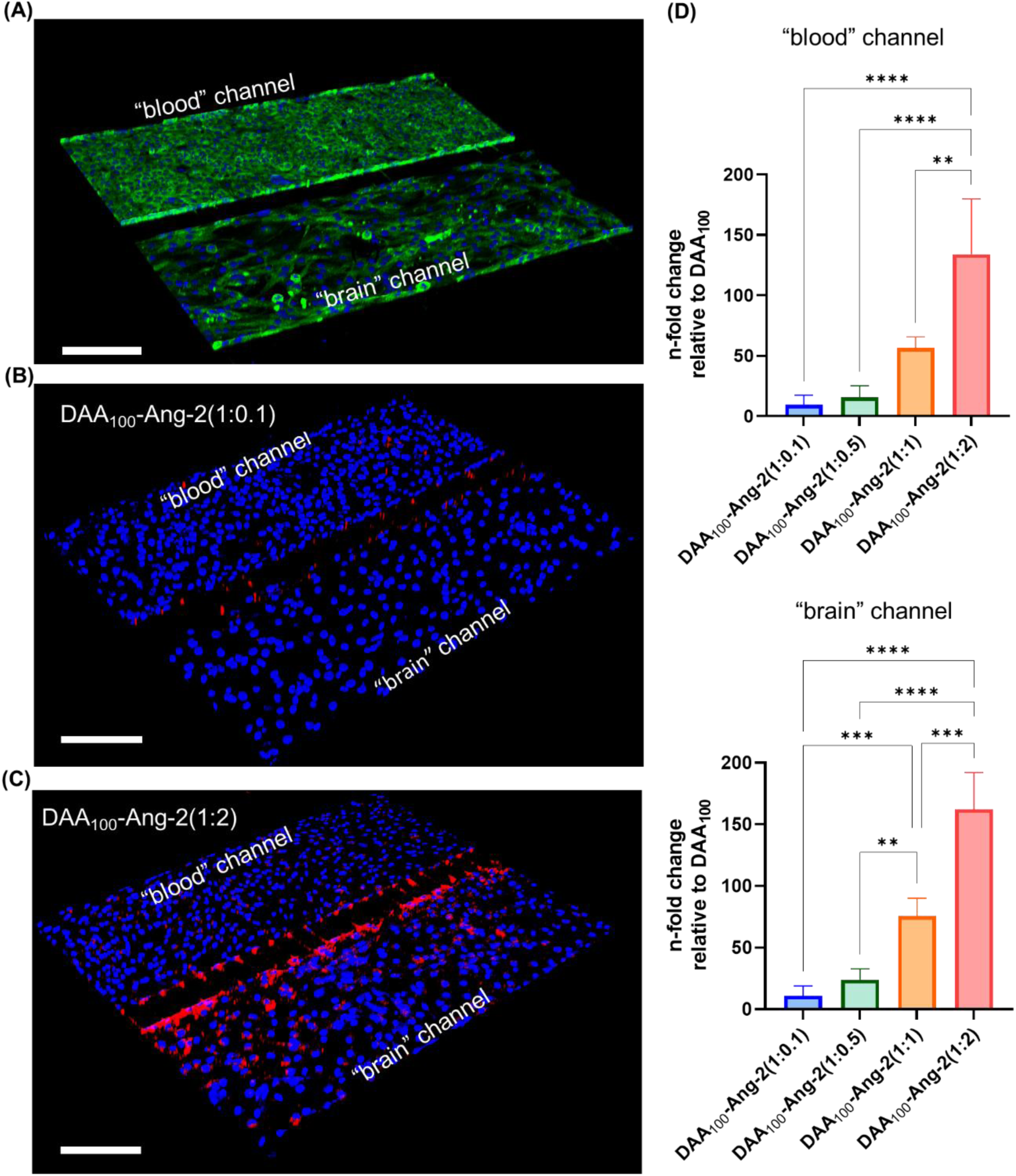
Evaluation of trafficking of Ang-2-conjugated nanoparticles in a microfluidic BBB-GBM-on-a-chip model. (A) Confocal fluorescence microscopy image of BBB-GBM-on-a-chip in 3D view (blue = nuclear, green = F-actin, scale bar = 200 µm. Representative images of BBB-GBM-on-a-chip treated with (B) DAA_100_-Ang-2(1:0.1) and (C) DAA_100_-Ang-2(1:2) (blue = nuclear, red = Cy5 in nanoparticles, scale bar = 200 µm). (D) Fold change of Ang-2 conjugated nanoparticles signal relative to control nanoparticles (DAA_100_) in “blood” and “brain” channels. One-way ANOVA test used with Tukey’s multiple comparisons test, * p < 0.05, ** p < 0.01, *** p < 0.001, **** p < 0.0001. The values are shown as mean ± SD (n = 3).

As the Ang-2 peptide in our designed nanoparticles primarily targets the LRP1 receptor, we further confirmed the expression and distribution of LRP1 in the BBB-GBM-on-a-chip using immunostaining. As demonstrated in **Figure S12**, U87 GBM cells (brain channel) have presented higher LRP1 expression compared to hCMEC/D3 cells (blood channel). This indicates that nanoparticles with a higher Ang-2 density may not only foster increased interaction with hCMEC/D3 cells in the blood channel but also enhance targeting and uptake by GBM cells in the brain channel, thereby further facilitating the differential BBB penetration and GBM uptake among nanoparticle variants. In addition, fluidic conditions have been shown to upregulate certain transmembrane proteins (e.g. caveolin-1 and claudin-11) in endothelial cells, potentially promoting a higher uptake and transcytosis of selective nanoparticles compared to static conditions^56, 58, 59^.

In previous studies, fluid flow has been demonstrated to change the absorption and selectivity of nanoparticles to receptors^60, 61^. Although our microfluidic model does not mimic all the complexities of the human vasculature, it does incorporate fluid flow and fluid shear stress. These factors might explain the observed differences in nanoparticle association and consequent transcytosis between Transwell and BBB-GBM-on-a-chip assays^26, 62^. For instance, in a static environment (e.g. Transwell assay), nanoparticles with a high density of Ang-2 are likely to have a higher avidity to LRP1, leading to strong membrane association or lysosomal degradation rather than transcytosis^23^. However, under fluid flow (e.g. microfluidic model), the fluid shear stress can diminish the avidity of Ang-2 toward LRP1, favoring higher transcytosis for nanoparticles with more Ang-2^63, 64^. Such a phenomenon was also reported for the transferrin receptor^65, 66^ where low-affinity transferrin receptor antibodies were transcytosed while high-affinity transferrin receptor antibodies were directed to lysosomes for degradation^47^.

### 2.7. Nanoparticle brain uptake study in mice

An overall safety evaluation of nanoparticles in mice was conducted before undertaking the brain uptake and biodistribution studies. Here, we confirmed the safety profiles of all nanoparticles using complete blood counting (CBC), alanine aminotransferase (ALT) and aspartate aminotransferase (AST) biochemical testing, and histology analyses of major body organs **(Figure S13 & S14)**.

The uptake of intravenously injected nanoparticles into the brains of Swiss-outbred mice was tested using *ex vivo* fluorescence scanning 24 h after administration. We chose to test the brain uptake of DAA100, DAA100-Ang-2(1:0.1), and DAA100-Ang-2(1:2), since these two Ang-2-conjugated formulations presented the most contradictory results in the previous *in vitro* Transwell and BBB-GBM-on-a-chip models. This would provide insight into the effect of Ang-2 conjugation and its surface density on the subsequent nanoparticle uptake into the brain. As shown in **Figure 6A**, DAA100-Ang-2(1:2) exhibited the highest nanoparticle fluorescence signals in the brains, of ∼ 6.8 times and 9.1 times higher than DAA100-Ang-2(1:0.1) and DAA100, respectively (**Figure 6B**). These notable differences in mouse brain accumulation between DAA100-Ang-2(1:2), DAA100-Ang-2(1:0.1), and DAA100 further confirmed the role of Ang-2 in enhancing BBB trafficking of nanoparticles. Conjugation of Ang-2 to the DAA100 nanoparticles did not result in a significant change in their biodistribution behaviour to the other major body organs **(Figure S15)**. All the tested nanoparticles presented strong fluorescence signals in lung, liver, and kidney, while lower signals were detected in the spleen and heart. This further suggested that the *in vivo* brain uptake seems to be dependent on the Ang-2 surface density. This was also reported for other Ang-2 polymeric nanoparticle systems^67^. Importantly, the *in vivo* result aligns better with the transport patterns observed in the BBB-GBM-on-a-chip rather than with the Transwell BBB model results. This reinforces the hypothesis that dynamic conditions could lead to different interactions of nanoparticles with cells and consequently affect the outcome of the preclinical *in vitro* assessments of nanoparticles.

**Figure 6.**
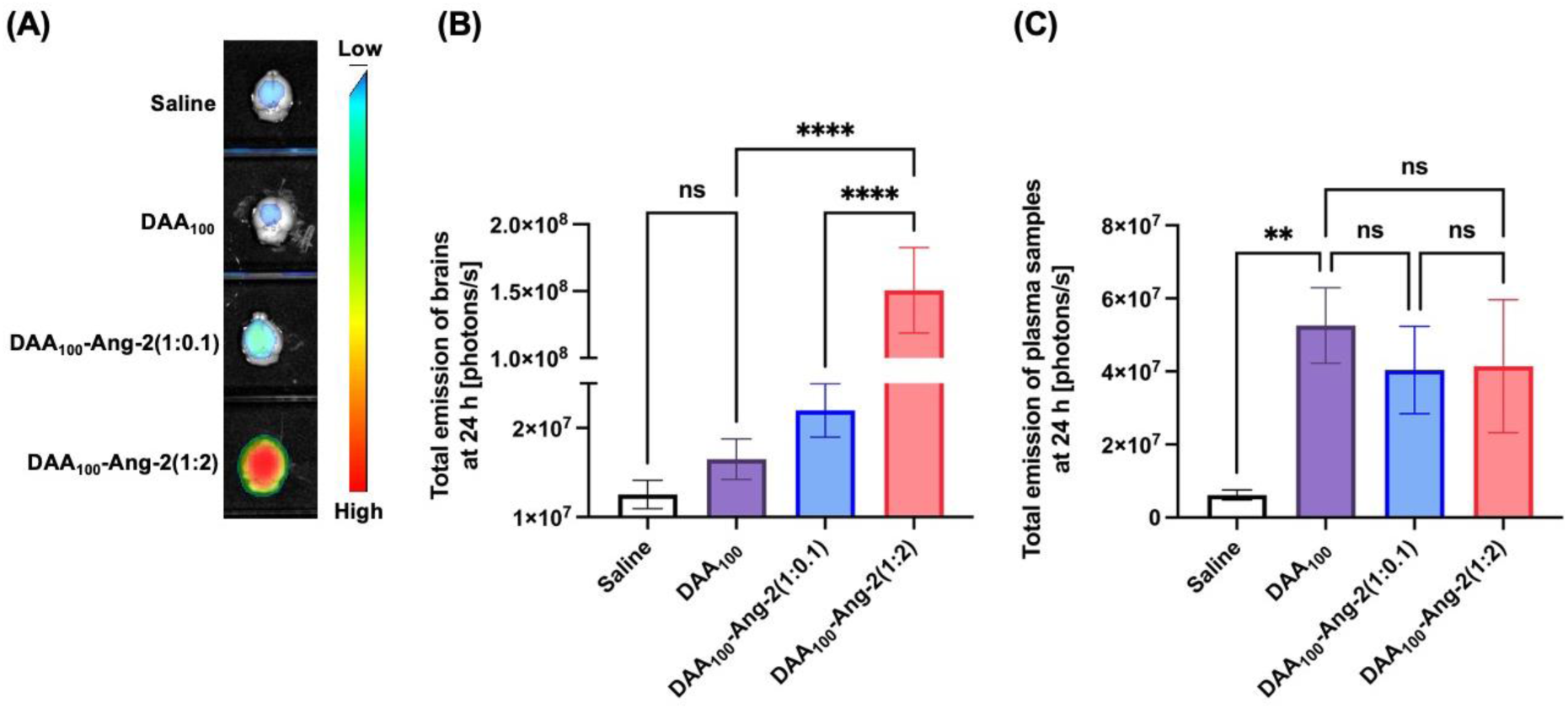
Brain accumulation of DAA_100_ with an increased molar ratio of Ang-2 at 24 h. (A) *Ex vivo* fluorescence photographs of whole brains imaged 24 h after intravenous injection of saline, DAA_100_, DAA_100_ -Ang-2(1:0.1), and DAA_100_-Ang-2(1:2). Total emissions [photons/s] of (B) brain samples and (C) diluted plasma samples at 24 h intravenous administration of different treatments. Data analysis was performed using One-way ANOVA test with Tukey’s multiple comparisons, * p < 0.05, ** p < 0.01, *** p < 0.001, **** p < 0.0001. The values are shown by mean ± SD, n = 3.

The BBB crossing ability of DAA100, DAA100 -Ang-2(1:0.1), and DAA100-Ang-2(1:2) in mice is less pronounced than in the microfluidic chips, which can be attributed to several *in vivo*-related factors. Firstly, some nanoparticles may be cleared by the immune system and accumulate in the spleen and liver^68, 69^. For instance, De Jong *et al.* reported that up to 46% of intravenously injected gold nanoparticles were located in the livers of rats within 24 h^70^. Additionally, the plasma protein adsorption *in vivo* has been found to mask the nanoparticle targeting surface, resulting in a reduction of specificity^71, 72^. Salvati *et al.* demonstrated that glycoprotein transferrin-coated silica nanoparticles lost their targetability to transferrin receptors after the formation of a protein corona^73^. Xiao *et al.* also found that nanoparticles coated with protein corona were less effective in penetrating BBB and targeting brain tumors *in vivo* than *in vitro*^74^. Such findings reveal that some potential constraints (e.g. absence of mononuclear phagocyte systems or a wide range of protein association) in the *in vitro* models may influence its nanoparticle assessment^75–77^.In summary, microfluidic organ-on-a-chip models like the BBB-GBM-on-a-chip, which mimic some aspects of the *in vivo* conditions, can offer a more precise and relevant evaluation of nanoparticles. In contrast, static Transwell might only provide limited information due to the lack of flow.

Despite the promising *in vivo* data, it should be acknowledged that *ex vivo* fluorescence scanning of brains does not account for vascular volume depletion which might result in overestimation of the nanoparticles accumulation in the brain parenchyma and represent a technical limitation of this methodology. The limited tissue penetration depth of the used excitation lasers represents another technical challenge. Radioisotopic assays could offer a more accurate and quantitative option in this regard, where uptake of radiolabelled-nanoparticles can be quantified using beta-scintillation counting of brain homogenates. A radiolabelled high molecular weight probe such as [^14^C] insulin can be used to calculate the brain vascular volume as previously reported^78^.

### 2.8. Application of nanoparticle system for doxorubicin delivery to GBM cells

We hypothesized that the Ang-2 conjugated nanoparticles could help chemotherapeutic drugs cross the BBB and target GBM cells for enhanced anti-proliferation effects. DAA100-Ang-2(1:2) was therefore selected for this study given its demonstrated high BBB penetration in both the microfluidic BBB-GBM-on-a-chip model and the *in vivo* study. For proof of concept, doxorubicin (Dox) was used as a chemotherapy drug. Dox is an effective anticancer drug showing cytotoxicity in many tumors, but it has limited ability to penetrate the BBB^79, 80^. The hydrophobic P(DAA-*co*-ALAM) core allowed for entrapment of the Dox via a Schiff-base reaction with the present ketones resulting in the formation of pH-responsive imine bonds. DAA100-Ang-2(1:2) presented an average Dox loading of 25.6 ± 3.4 µg per mg of nanoparticles, which was slightly higher than DAA100 (21.7 ± 0.1 µg/mg). The colloidal stability of nanoparticles after Dox loading was confirmed using DLS, where Dox@DAA100-Ang(1:2) and Dox@DAA100 showed average hydrodynamic diameters of 161.9 ± 24.4 nm and 100.2 ± 24.1 nm, respectively, with PDI of < 0.3. Dox release studies were conducted under conditions representing the endolysosomal (pH 5) and physiological microenvironment (pH 7.4). Both Dox-loaded nanoparticles showed pH-dependent release kinetics over the tested timeframe, with faster and more Dox release (up to 50-65% at 24 h) under acidic conditions (pH 5) **(Figure 7A and 7B)**. This confirmed that the designed pH-responsive imine bond could provide a cleavable Dox release under slightly acid conditions and potentially promote higher release in the acidic glioblastoma endosomes. The retained unreleased Dox could be attributed to interactions of the relatively hydrophobic molecule with the hydrophobic core of the nanoparticles.

**Figure 7.**
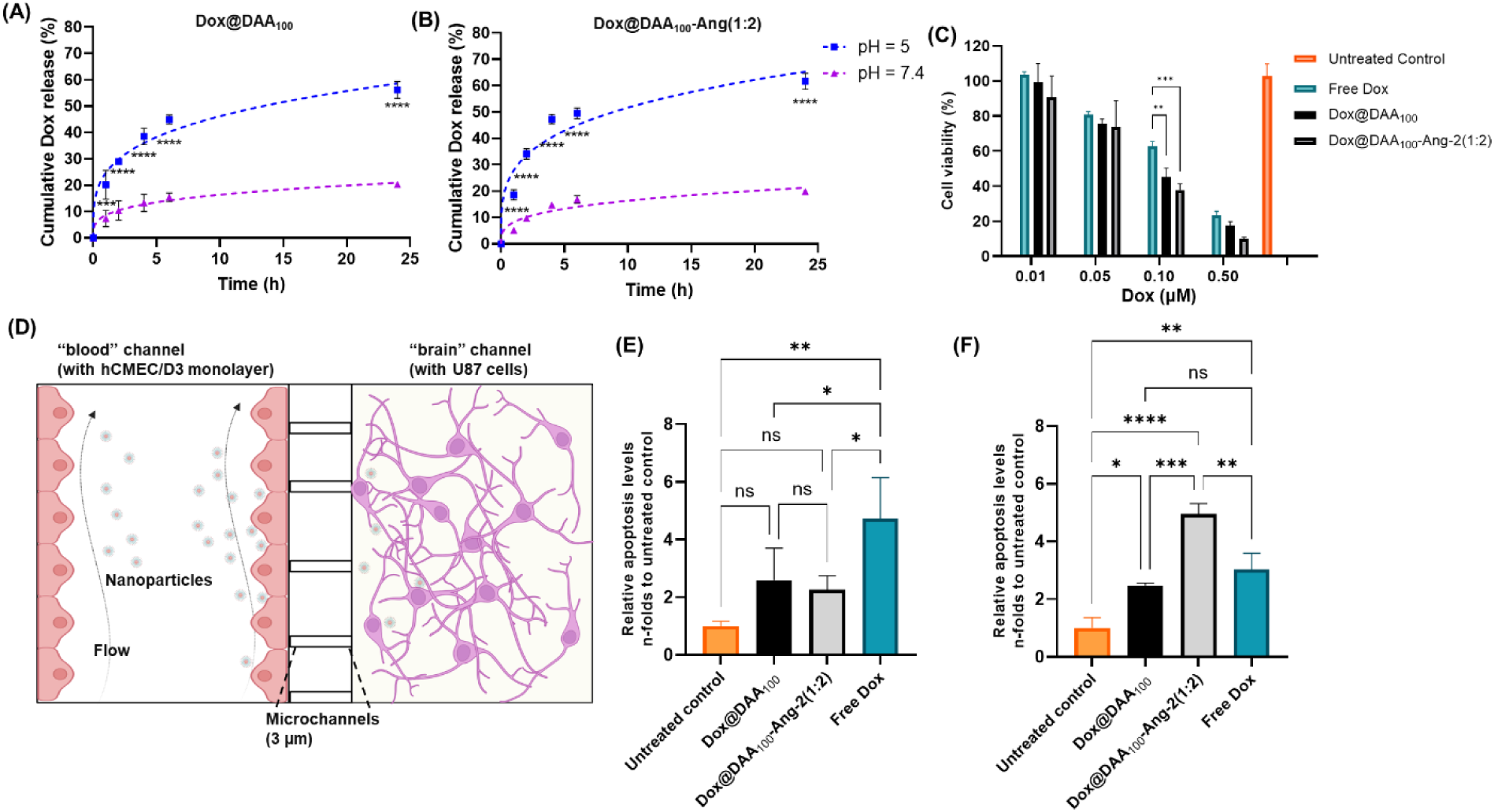
Evaluation of Dox-loaded nanoparticles against U87 GBM cells *in vitro*. *In vitro* drug release of Dox from (A) Dox@DAA_100_ and (B) Dox@DAA_100_-Ang-2(1:2) at pH 5 and pH 7.4, using the two-way ANOVA with Tukey’s multiple comparisons test, * p < 0.05, ** p < 0.01, *** p < 0.001, **** p < 0.0001, n = 3. (C) Cytotoxicity assessment of various concentrations of Free Dox, Dox@DAA_100_ and Dox@DAA_100_-Ang-2(1:2) (0.01, 0.05, 0.10, and 0.50 µM) on U87 after 72 h treatment, using the two-way ANOVA with Tukey’s multiple comparisons test, * p < 0.05, ** p < 0.01, *** p < 0.001, **** p < 0.0001, n = 3. All data are shown as mean ± SD, n = 3. (D) Schematic of the BBB-GBM-on-a-chip model consisting of two channels. The blood channel was formed by a monolayer of hCMEC/D3 cells over 3 µm microchannels under a fluidic environment. The brain channel was seeded by U87 cells receiving medium and Dox-loaded nanoparticles from the blood channel. Relative apoptosis levels (n-fold change to the untreated control chips) of activated caspase-3/7 (E) in “blood” channels and (F) in “brain” channels, respectively, using the one-way ANOVA with Tukey’s multiple comparisons test, * p < 0.05, ** p < 0.01, *** p < 0.001, **** p < 0.0001, n = 3. All data are shown as mean ± SD.

The *in vitro* cytotoxicity of Dox-loaded nanoparticles on GBM was first assessed using the U87 glioblastoma cell line in 2D cell culture. Dox@DAA100-Ang(1:2) and Dox@DAA100 displayed a higher U87 cell toxicity than free Dox at all tested concentrations. This could be attributed to the potential increased uptake of Dox in the nanoparticles compared to free Dox as well as the pumping out of free Dox by P-glycoprotein efflux transporters^81–83^. Furthermore, compared to the control nanoparticles, Dox@DAA100-Ang(1:2) showed higher cytotoxicity, which could be attributed to the higher specificity of the Ang-2 coating (**Figure 7C**).

The designed microfluidic BBB-GBM-on-a-chip was further used to assess the anticancer effect of the Dox-loaded Ang-2 conjugated nanoparticles. Firstly, to ensure the Dox concentration did not affect the integrity of the BBB model, FITC-dextran (25 µg/mL, 10 kDa) was co-administered with different concentrations of Dox into the blood channels of the chips under flow conditions. Permeability of FITC-dextran through chips treated with Dox at the concentration of 1 µM was not significantly different from FITC-dextran in non-Dox treated chips suggesting an optimal concentration range for the next investigations (**Figure S16**).

The anticancer effect of nanoparticles in the chips was assessed using a caspase-3/7 assay (caspase 3 and caspase 7 are proteins that are both activated universally during apoptosis and can be stained by a DNA-binding dye ^84^. After nanoparticle treatment, cells in both “blood” channels and “brain” channels were stained using a caspase-3/7 staining kit, and the fluorescence intensities of caspase-3/7 activated cells in both channels were compared to the signals from the untreated control, representing relative apoptosis levels. In the “blood” channels, cells treated with Dox@DAA100-Ang-2(1:2) shared a similar caspase-3/7 activation to the Dox@DAA100, both being lower than the free Dox (**Figure 7E**). In contrast, in the “brain” channels, cells treated with Dox@DAA100-Ang-2(1:2) displayed 100% and 60% higher activated caspase-3/7 levels compared to the cells treated with Dox@DAA100 and free Dox, respectively, which was consistent to our results in 2D cell assay (**Figure 7F**). These results demonstrate that Ang-2 conjugated nanoparticles can not only promote better BBB penetration but can also enhance the anticancer effect of Dox to U87 GBM cells. In addition, although free Dox shared a similar level of apoptosis in GBM cells with Dox@DAA100 in the “brain” channels, free Dox triggered a higher toxicity in the BBB compared to Dox@DAA100, as indicated in the “blood” channels. This suggests that free Dox might trigger some disruptive pathways in BBB, leading to an increased uptake in the brain channel. For instance, Lopez-Ramirez *et al.* found that activation of specific caspase 3 might trigger cytokine-induced disruption of tight and adherent junctions in the BBB^85^.

In summary, the BBB-GBM-on-a-chip model enabled new insights into the cytotoxicity of nanoparticles or drugs when crossing the BBB and combating GBM in a dynamic environment. In the BBB, Dox-loaded nanoparticles exhibited a high degree of apoptosis and still demonstrated reduced apoptosis levels than the free Dox. In addition, Dox@DAA100-Ang-2(1:2) showed a higher anticancer effect against GBM cells compared to both free Dox and Dox@DAA100-Ang-2(1:2), suggesting the advantages of the designed Ang-2 conjugated polymeric system in helping Dox traverse the BBB and target GBM.

To that end, our future studies will explore the use of an optimized formulation for anticancer delivery to GBM in an orthotopic brain tumor model. Radiolabeled nanoparticles will be potentially used for a more quantitative tracking of brain delivery.

## 4. Conclusion

In this study, we synthesized a multifunctional pH-responsive polymeric nanoparticle and tailored its surface with various Ang-2 densities. This diverse Ang-2 nanoparticle library was systematically employed to investigate the impact of Ang-2 surface density on BBB penetration through a series of increasingly complex *in vitro* models and an *in vivo* study. Our findings revealed a significantly different interplay between Ang-2 surface density and BBB penetration for the different models. In 2D cell culture, a positive correlation between Ang-2 surface density and hCMEC/D3 cellular association was observed with a distinctive inflection range (i.e., DAA100-Ang-2(1:2)). Interestingly, non-primary cell uptake pathways such as caveolae-mediated endocytosis and micropinocytosis played a more prominent role in nanoparticles with a higher Ang-2 density. However, in the Transwell model, higher Ang-2 surface density correlated negatively with BBB transportation, with the highest Ang density resulting in the lowest BBB penetration. In contrast, the BBB-GBM-on-a-chip demonstrated an opposite trend, with higher Ang-2 densities leading to increased nanoparticle BBB transport. These contrasting results may be attributed to variances in avidity in static versus fluidic conditions, where dynamic environments could reduce the avidity of nanoparticles due to flow forces limiting endolysosomal degradation and sorting. The *in vivo* model supported the findings of the BBB-GBM-on-a-chip, further highlighting the crucial role of flow in affecting avidity and BBB transportation of nanoparticles. Finally, Dox was successfully loaded into the polymeric nanoparticles with optimal Ang-2 density (1:2), showing a controllable pH-responsive release while maintaining colloidal stability. The enhanced anticancer effects against U87 GBM cells were demonstrated in a 2D cell assay and the BBB-GBM-on-a-chip model. More insights and translational potential may be gleaned from *in vivo* GBM orthotopic models, further validating the nanoparticles and the relevance of the BBB-GBM-on-a-chip model for future studies.

## 5. Experimental Section

### 5.1. Materials

The CellTiter-Glo® luminescence cell viability assay kit was purchased from Promega (Sydney Corporate Park, Australia). Cyanine5 amine (Cy5) and Cyanine3 DBCO(Cy3-DBCO) were purchased from Lumiprobe. (2,2’-Azobis[2-(2-imidazolin-2-yl)propane]dihydrochloride (VAZO-44) initiator was purchased from Fujifilm (Minato City, Japan). 2,2′-Azobis(2-methylpropionitrile) (AIBN), diacetone acrylamide (DAA), dimethylacrylamide (DMA), allylamine, acryloyl chloride, 2-(dodecylthiocarbonothioylthio)-2-methylpropionic acid 3-azido-1-propanol ester chain transfer agent (CTA-azide) were purchased from Sigma-Aldrich (Macquarie Park, Australia). DBCO-modified Ang-2 peptide (TFFYGGSRGKRNNFKTEEYC(Mal-DBCO)) was purchased from ChinaPeptide (Suzhou, China). Endothelial cell basal medium and endothelial cell growth factor kit were purchased from Lonza (Mount Waverley, Australia). Water (HPLC grade) was obtained with a Milli-Q Advantage A10 water purification system (Merck Millipore, Bayswater, Australia). All other chemical reagents, solvents, and mediums were purchased from Sigma-Aldrich (Macquarie Park, Australia) unless stated otherwise.

### 5.2. Characterization methods

Molecular weight properties of the polymers were analyzed by gel permeation chromatography (SEC) performed on a Shimadzu system equipped with a CMB-20A controller system, an SIL-20A HT autosampler, an LC-20AT tandem pump system, a DGU-20A degasser unit, a CTO-20AC column oven, a RDI-10A refractive index detector, and 4× Waters Styragel columns (HT2, HT3, HT4, and HT5, each 300 mm × 7.8 mm^2^, providing an effective molar mass range from 100 to 4 × 10^6^). *N*,*N*-Dimethylacetamide (DMAc) (containing 4.34 g/L lithium bromide (LiBr)) was used as an eluent with a flow rate of 1 mL/min at 80 °C. Number (Mn) and weight-average (Mw) molar masses were evaluated using Shimadzu LC Solution software. The SEC columns were calibrated with low-dispersity poly(methyl methacrylate) standards (Polymer Laboratories) ranging from 3100 to 650,000 g/mol, and molar masses are reported as poly(methyl methacrylate) equivalents. A third-order polynomial was used to fit the log Mp *vs* time calibration curve, which was near linear across the molar mass ranges. Aqueous gel permeation chromatography was performed on a GE Healthcare ÄKTA pure purification system. A Superdex 200 Increase 10/300 GL column was utilized for the hydrodynamic analysis of protein/polymer samples at a flow rate of 0.5 mL/min with PBS as the mobile phase (pH 7.4).

^1^H NMR spectra were recorded on a Varian UNITY-plus 400 NMR spectrometer using CDCl3, Methanol-*d*4, and D2O as the solvent. Liquid chromatography-mass spectrometry (LC-MS) data were acquired on a Waters e2695 HPLC equipped with a 2998 photodiode array (PDA) detector and an ACQUITY QDa mass detector. Ultraviolet-visible spectrophotometry (UV-Vis) was recorded using a UV-2450PC spectrophotometer from Shimadzu (Japan).

Transmission electron microscopy (TEM) images were used to measure the nanoparticle size and visualize morphological traits. TEM samples were prepared via drop casting the suspended sample of interest (in Milli-Q water or PBS) onto copper grids. Uranyl acetate staining was applied, and the samples were dried. The samples were examined using a Philips Tecnai 12 TEM at an operating voltage of 120 kV. Images were recorded using a FEI Eagle 4k × 4k CCD camera. Low dose conditions (<10 e^-^/Å^2^) were used to avoid damage to samples.

Dynamic light scattering (DLS) was performed on a Malvern Instruments Zetasizer Nano instrument ZEN3600 with a 4 mW 633 nm HeNe gas laser. An Avalanche photodiode detector measured the backscatter light at an angle of 173° relative to the angle of the incident light beam. Samples were dissolved in Milli-Q water or PBS (pH 7.4). A single-use folded capillary cell commercialized by Malvern Instruments (model DTS 1070) was used for measuring surface zeta potential.

### 5.3. Synthesis of allyl acrylamide (ALAM)

The synthesis of ALAM was described previously by Qu *et al.*^30^. Briefly, allylamine (0.92 g, 9.7 mmol) and triethylamine (Et3N) (2.0 g, 0.75 mmol) were combined and dissolved in dichloromethane (DCM) (12.5 mL) in an ice bath, followed by a dropwise addition of acryloyl chloride (0.94 g, 10 mmol). The reaction mixture was stirred in an ice bath for 1 h before moving to room temperature (RT) for 24 h. The product was retrieved by three washes with 0.1 M HCl and then brine followed by drying over MgSO4.

### 5.4. Synthesis of Cy5-acrylamide

The synthesis procedure of Cy5-acrylamide was similar to the ALAM synthesis. Briefly, Cy5-amine (5.0 mg, 8.6 µmol) and Et3N (2.6 mg, 25 µmol) were combined and dissolved in DCM (1 mL) in an ice bath by stirring, followed by a dropwise addition of acryloyl chloride (0.93 mg, 0.010 mmol). The reaction mixture was reacted in an ice bath for 1 h before moving to RT for 24 h. The product was obtained by three washes with 0.1 M HCl, followed by brine, and dried over by MgSO4. The purified product was then characterized using liquid chromatography-mass spectrometry (LC-MS).

### 5.5. Synthesis of poly(dimethylacrylamide-*co*-Cy5-acrylamide)(P(DMA-*co*-Cy5)) macro-CTA

2-(Dodecylthio-carbonothioylthio)-2-methylpropionic acid 3-azido-1-propanol ester (CTA-azide) (6.7 mg, 15 µmol), Cy5-acrylamide (3.4 mg, 3.8 µmol), AIBN (20 µg, 1.5 µmol) and dimethylacrylamide (74 mg, 0.25 mmol) were dissolved in dioxane (0.5 mL). The mixed solution was sparged with nitrogen gas in an ice bath for 30 min and the polymerization reaction was conducted at 70 °C 24 h. The product was then purified three times by washing with diethyl ether and characterized by ^1^H NMR and SEC.

### 5.6. Synthesis of poly(dimethylacrylamide-*co*-Cy5-acrylamide)-block-poly(diacetone acrylamide-co-allyl acrylamide) (P(DMA-*co*-Cy5)-*b*-P(DAA-*co*-ALAM)) block copolymer nanoparticles

P(DMA-*co*-Cy5)-*b*-P(DAA-*co*-ALAM) block copolymer nanoparticles were synthesized using aqueous RAFT dispersion polymerization with a molar feed ratio of DAA:ALAM:P(DMA-*co*-Cy5):VAZO-44 = 100:1:1:0.25. P(DMA-*co*-Cy5) as a macro-CTA (5.0 mg, 1.1 µmol), DAA (18 mg, 110 µmol), ALAM (0.24 mg, 1.1 µmol), and VAZO-44(90 µg, 0.28 µmol) were dissolved in 0.4 mL Milli-Q water. The reaction vessel was sealed and gently sparged with nitrogen gas for 30 min in an ice bath. The reaction was then immersed into a preheated oil bath at 70 °C for 24 h. Afterward, the polymerization was terminated by cooling down to RT for 5 min and exposed to air. The obtained nanoparticle formulations were then purified by dialysis (molecular weight cut-off (MWCO):10 KDa) against Milli-Q water (two solvent changes) and finally characterized by SEC, ^1^H NMR, DLS, TEM and Nanosight (PARTICLE METRIX, PMX120).

### 5.7. Synthesis of Ang-2-conjugated P(DMA-co-Cy5)-b-P(DAA-co-ALAM) copolymer nanoparticles

Nanoparticles with different Ang-2 density conjugations were prepared via copper-free click reaction using different reaction molar ratios of the polymer P(DMA-*co*-Cy5)-*b*-P(DAA-*co*-ALAM) to Ang-2-DBCO. Briefly, P(DMA-*co*-Cy5)-*b*-P(DAA-*co*-ALAM) and Ang-2 were mixed in four feeding molar ratios (1:0.1, 1:0.5, 1:1, and 1:2). P(DMA-co-Cy5)-b-P(DAA-co-ALAM) was maintained at the concentration of 3.3 mg/mL in 0.3 mL Milli-Q water for each sample of preparation. These mixtures were gently vortexed for 10 min before shaking overnight at RT. Afterward, the conjugated samples were purified by dialysis membranes of MWCO 10 KDa against Milli-Q water (two solvent changes). The conjugation of Ang-2 in these purified products was determined using a micro-BCA assay (Thermo Scientific, Australia). We also used Cyanine3-DBCO (Cy3-DBCO) to indirectly determine the number of Ang-2 per polymer. Briefly, Cy3-DBCO (0.44 mg/mL) and Ang-2 conjugated/unconjugated nanoparticles (5 mg/mL) were mixed in 0.1 mL of DMSO before vertexing for 10 min and shaking 48 hours at RT. Afterward, the Cy3 conjugated samples were purified by dialysis membranes of MWCO 10 KDa against 80 % of Ethanol in water (two solvent changes) for 72 hours. The amount of conjugated Cy3-DBCO, representing unreacted azido groups, was determined by a Cy3-DBCO standard curve.

### 5.8. Doxorubicin loading and release kinetics

Dox (0.8 mg) was dissolved in 160 µL of PBS (pH 8) and dropwise added to 1 mg of synthesized nanoparticles (340 µL in PBS pH 8). The mixture was sonicated for 20 s and vortexed for 10 min before stirring at RT overnight. The reaction mixture was then purified by dialysis (MWCO: 10KDa) against Milli-Q water (two solvent changes). The amount of Dox loading was determined by UV–vis spectrophotometry (UV-2450PC, Shimadzu, Japan) at the absorbance of 496 nm, according to the standard curve obtained from a dilution series of Dox in Milli-Q water.

*In vitro* release profiles of Dox from polymer nanoparticles were investigated in PBS (pH 7.4) and MES buffer (pH 5). The Dox-loaded nanoparticles (0.5 mg) were suspended in PBS (0.5 mL) and introduced into a dialysis bag (MWCO: 10KDa). The release experiment was initiated by placing the end-sealed bag in 2 mL of PBS at 37 °C with continuous shaking at 100 rpm. At selected intervals (1 h, 2 h, 4 h, 6 h, and 24 h), 0.2 mL of buffer was collected for measurement using a PerkinElmer EnSpire multimode plate reader (excitation: 472 nm and emission: 589 nm) and an equal volume of fresh buffer was replenished. The amount of released Dox was determined by extrapolation against a standard curve obtained from a dilution series of Dox in Milli-Q water.

### 5.9. Cell culture

Immortalized human brain microvascular endothelial cells (hCMEC/D3) cells and human glioblastoma U87 cells were purchased from Merck. The hCMEC/D3 cell line was maintained using EC growth basal medium-2 (EBM-2, Lonza) with supplements of 5% FBS and growth factors as reported^86^. hCMEC/D3 cells were cultured on collagen-coated (150 µg/mL) T-75 tissue culture flask at 37 °C in 5% CO2 in the passages of 28 to 35 according to the supplier’s protocol. The medium was changed every 2 to 3 days before reaching confluency. U87 cells were maintained in Dulbecco’s Modified Eagle Medium (DMEM) with supplements of 10% FBS and 1% antibiotic–antimycotic (Life Technologies, 15240062). U87 cells were cultured on a culture flask at 37 °C in 5% CO2 and the medium was changed every 3 to 4 days before reaching confluency.

### 5.10. Cytocompatibility assay

The biocompatibility of polymeric nanoparticles on hCMEC/D3 and U87 cell lines was determined using a CellTiter-Glo® luminescence cell viability assay (Promega). Briefly, hCMEC/D3 cells and U87 cells were seeded into a 96-well white polystyrene microplate (Sigma) at a density of 10,000 and 5,000 cells per well, respectively. Cells were allowed to grow in their respective cell culture medium for 1 day. Subsequently, cells were treated with different concentrations (10, 50, 100, and 200 µg/mL) of polymeric nanoparticles. Cells without any treatment were used as controls, and each experiment was carried out in triplicate. After incubating cells for 48 h, a CellTiter-Glo® luminescence cell viability assay was used to evaluate the cell viability. Briefly, 100 µL of the solution of the assay kit was added to 100 µL of the medium in each well, and the plates were gently shaken at RT for 15 min. Finally, the luminescence intensity of each sample was obtained using a PerkinElmer EnSpire multimode plate reader.

### 5.11. Cellular association of polymeric nanoparticles via flow cytometry and confocal microscopy

Flow cytometry was used to confirm the cellular uptake of polymeric nanoparticles in hCMEC/D3. Briefly, hCMEC/D3 cells were seeded into a 24-well plate at the density of 12 × 10^4^ cells per well, respectively, and cultured overnight. The cells were then washed twice in PBS and incubated with 10 µg/mL of Cy5-labelled polymeric nanoparticles for 1 h at 37 °C supplied with 5% CO2. Afterward, the cells were washed three times with PBS to remove any unattached nanoparticles and harvested by trypsin and centrifugation for 3 min at 180 rcf. The resulting cell pellets were dispersed in cold PBS and stained with PI for 5 min to assess cell viability. Samples were then analyzed using flow cytometry (BD FACS Canto II) for Cy5 and PI fluorescence signals. Cellular association percentage was calculated as the number of cells that displayed positive fluorescence signals compared to untreated cells (Cy5 negative).

For visualizing the cellular association and uptake, confocal microscopy was used. hCMEC/D3 and U87 cells were seeded into an 8-well chamber glass coverslip (Ibidi) at the density of 1 × 10^5^ cells and 5 × 10^4^ cells per well, respectively, and allowed to attach and grow for 24 h. The cells were washed twice in PBS and then treated with Cy5-labelled polymeric nanoparticles (10 µg/mL). After 1 h, the wells were washed twice in PBS and fixed using 4% paraformaldehyde (PFA) for 15 min at RT followed by permeabilization with 0.1% Triton-X-100 in PBS for 5 min at RT. The cells were then washed by PBS twice and incubated with Hoechst 33342 (1:2000 Thermo Fisher Scientific) and Rhodamine Phalloidin (1:100 Thermo Fisher Scientific) for 30 min. Afterward, the cells were washed three times with PBS, and the images were taken using confocal microscopy (Leica TCS SP8, Leica Microsystem). Fluorophores were excited by 405, 561, and 647 nm laser lines. The laser powers and gains for every channel were adjusted against untreated controls to avoid the detection of sample autofluorescence.

### 5.12. Nanoparticles uptake mechanism in hCMEC/D3 cells

Three pharmacological blockers of different endocytic pathways were used to investigate the mechanisms involved in the uptake of nanoparticles by hCMEC/D3 cells. hCMEC/D3 cells were seeded in a 24-well plate at the density of 12 ×10^4^ cells/well. After 24 h, cells were pre-treated with blockers, nystatin (50 µg/mL, Sigma, N4014), chlorpromazine (15 µg/mL, Sigma, C8138), and amiloride hydrochloride hydrate (125 µg/mL, Sigma, A7410) for 1 h. Afterward, the blockers were washed off and cells were incubated with nanoparticles of different Ang-2 densities (10 µg/mL) in fresh media for 1 h. The cells were then washed thrice with PBS to remove any unattached nanoparticles and the cells were harvested by trypsin and centrifugation for 3 min at 180 rcf. The resulting cell pellets were dispersed in cold PBS and stained with PI (5 µg/mL) for 5 min to assess cell viability. Samples were then analyzed using flow cytometry (BD FACS Canto II) for Cy5 and PI fluorescence signals. Cellular association percentage was calculated as the number of cells that displayed fluorescence signals compared to untreated cells (Cy5 negative).

### 5.13. BBB Transwell model preparation, and nanoparticles assessment

The BBB Transwell model was established as reported^28^. The transendothelial electrical resistance (TEER) was assessed every day for 7 to 10 days using a Millicell ERS-2 voltammeter EVOM2 and an STX02 chopstick electrode (Merck Millipore, Bayswater Australia). The TEER value was calculated by subtracting the resistance of a blank insert from an insert containing a monolayer and then multiplying it by the surface area of the insert.

For assessment of nanoparticle BBB penetration in the Transwell model, on day 7 of monolayer culturing and once an acceptable TEER value was achieved, Ang-2-conjugated polymeric nanoparticles (200 µL, 200 µg/mL) were incubated at the donor chamber in complete culture medium and the receptor chamber was filled with 600 µL of complete culture media and further incubated at 37 °C and 5% CO2 for 48 h. After that, a 200 µL aliquot of multiple samples was collected from the receptor chamber and measured for the fluorescence intensity (assessing the Cy5 signal) by using a PerkinElmer EnSpire multimode plate reader. The concentration was determined using a calibration curve of different concentrations of nanoparticles in the cell medium. The percentage of nanoparticles in the receptor chamber was calculated as the fluorescence intensity from the receptor chamber was divided by the original fluorescence intensity of nanoparticles that were placed in the donor chamber. The apparent permeability coefficients (*Papp*), as the indication of the nanoparticle’s permeation, were calculated according to the following equation^87, 88^:

Where Vr (mL) is the volume of the receptor chamber, dC/dt is the slope of the cumulative concentration of the nanoparticles in the receptor chamber over time, A (cm^2^) is the surface area of the inset, and C (µg/mL) is the initial concentration of nanoparticles that was placed into the donor chamber^27^.

### 5.14. Cytotoxicity of Dox-loaded nanoparticles against U87 cells in 2D cell culture plate

The anticancer effect of the Dox-loaded nanoparticles in U87 cells was evaluated using CellTiter-Glo® luminescence cell viability assay (Promega). Briefly, U87 cells were seeded into a 96-well white polystyrene microplate at 2500 cells/well and incubated for 24 h at 37 °C in a humidified 5% CO2 incubator. Cells were treated for 72 h with either free Dox, Dox@DAA100, or Dox@DAA100-Ang-2(1:2) using equivalent Dox concentrations of 0.01, 0.05, 0.1, and 0.5 µM diluted in DMEM. Cells without treatment were used as the control. After 72 h, cell viability was demonstrated using the same aforementioned method under section 5.10.

### 5.15. BBB-GBM-on-a-chip model establishment, LRP1 immunostaining, nanoparticles permeability, and Dox loaded nanoparticles assessment

The BBB-GBM-on-a-chip model was established following our previously published method with some modifications^28^. The device features 8 subunits, and each subunit is comprised of three parallel main channels: blood, brain, and medium channel. Each channel is 500 µm wide, 100 µm height, and 2 cm length. An array of 3 µm width, 80 µm length, and 5 µm height of microchannels connects each blood channel with its adjacent brain channel. The blood channel serves as a simplified version of the BBB by establishing hCMEC/D3 monolayer, while the brain channel allowed us to assess nanoparticle penetration by culturing U87 cells.

hCMEC/D3 cells were seeded to the blood channel in the concentration of 8 x 10^6^ cells/mL and cultured for three days. Matrigel (Corning) was then added to the brain channel to facilitate cell cell-attached environment. Afterward, U87 cells were seeded to the brain channel in the concentration of 4 x 10^6^ cells/mL. Both hCMEC/D3 and U87 cells were further cultured for 2 days before conducting LRP1 immunostaining and nanoparticle testing.

For LRP1 immunostaining, cells in the chips (brain and blood channels) were fixed using 4% PFA and washed with PBS three times after 10 min. Afterward, cells were permeabilized with 0.1% Triton-X for 10 min. Then, a blocking buffer of 3 % goat serum (BSA) was used to block the cells for 1 h at room temperature. After 1 h, the cells were washed with PBS and incubated with rabbit anti-LRP1 antibody in blocking buffer (1:200) overnight at 4°C. Afterward, cells were washed with PBS and incubated for 1 h at room temperature with goat anti-Rabbit IgG (H+L) highly cross-adsorbed secondary antibody, Alexa Fluor™ 546 in a blocking buffer (1:200). Finally, cells were incubated with Hoechst 33342 (1:1000) for 5 min and washed with PBS three times before imaging using a Leica SP-8 Lightning confocal microscope.

For the nanoparticle testing, nanoparticles were diluted in EBM-2 medium to the concentration of 40 µg/mL and flowed through the blood channel at the flow rate of 0.25 µL/min for 24 h using a programmable syringe pump (Harvard Apparatus, Inc.). Afterward, the chips were detached from the syringe pump and carefully rinsed with PBS to remove any unbonded nanoparticles. Then, cells in both channels were fixed with 4% PFA, stained with DAPI, and examined under a confocal microscope (Leica TCS SP8, Leica Microsystems, Macquarie Park, Australia).

For the Dox loaded nanoparticles assessment, an optimal Dox concentration that did not interrupt the integrity of BBB in the chips during cytotoxic assessment was determined using FITC-dextran (10 kDa), a fluorescent probe with limited permeability across BBB. Briefly, FITC-dextran (25 µg/mL) was co-administered with free Dox at concentrations of 0.5, 1, and 2 µM into the blood channels at a flow rate of 0.5 µL/min for 2 h using a programmable syringe pump (Harvard Apparatus, Inc.). The blank chips (positive control, without cell seeded) and cell-seeded chips (negative control) were flown with 10 kDa FTTC-dextran (25 µg/mL) in the same flow rate and time mentioned above. Afterward, all chips were imaged using confocal microscopy, and the relative fluorescence intensity (RFU) of crossed FITC-dextran in the brain channel was measured using ImageJ (NIH, Bethesda, MD).

For Dox-loaded nanoparticles assessment, Dox@DAA100, Dox@DAA100-Ang-2(1:2), and free Dox (equivalent to 1μM of Dox) were flown through the blood channel in the BBB-GBM chips at a flow rate of 0.5 µL/min for 2 h using a programmable syringe pump (Harvard Apparatus, Inc.). Afterward, blood channels were changed to fresh medium and incubated for a further 48 h at 37 °C in a cell incubator supplied with 5% CO2. Cells in both blood and brain channels were incubated with Hoechst 33342 (20 µM Thermo Fisher Scientific) and CellEvent™ caspase-3/7 green detection reagent (2 µM Thermo Fisher Scientific) for 1 h and then subjected to confocal microscopy imaging (Leica TCS SP8, Leica Microsystems). The relative fluorescence intensity of caspase-3/7 from two channels of the chips was measured using ImageJ (NIH, Bethesda, MD).

### 5.16. *In vivo* brain accumulation of nanoparticles

This animal experiment was approved by the Monash Institute of Pharmaceutical Sciences Animal Ethics Committee (MIPS 36203) and performed in accordance with the Guidelines for the Care and Use of Animals for Scientific Purposes (National Health and Medical Research Council). Male Swiss-outbred mice (6 - 8 weeks of age) were injected with different formulations of nanoparticles, DAA100, DAA100-Ang-2 (1:0.1), and DAA100-Ang-2 (1:2), at doses of 10 mg/kg using tail vein injection, and saline was used as the negative control. After 24 h, mice were anesthetized using isoflurane inhalation, and a cardiac puncture was performed to collect blood samples. The mice were then humanely killed, and the brain samples were collected for *ex vivo* analysis. Blood samples were centrifuged at 1500 g for 10 min and plasma samples were separated. Both plasma samples and whole brains were scanned with an optical imaging system (Ami HT, Spectral Instruments Imaging) using 605/670 nm excitation/emission wavelength. Images were analyzed using Aura Software.

### 5.17. Hematoxylin & eosin staining and blood biochemical study

This animal experiment was approved by the Animal Health and Use Committee of Northwestern Polytechnical University (2024JC-YBQN-0919).study and performed following the guidelines for the Care and Use of Laboratory Animals of the Chinese Animal Welfare Committee. Male Swiss-outbred mice (6 - 8 weeks of age) were injected with all formulations of nanoparticles at doses of 10 mg/kg using tail vein injection, and saline was used as the negative control. After 24 h, mice were anesthetized using isoflurane inhalation, and a cardiac puncture was performed to collect blood samples. An automated hematology analyzer and biochemical analyzer were used to determine white blood cells, lymphocyte, granulocyte, platelet, hemoglobin, alanine aminotransferase and aspartate aminotransferase. The mice were humanely killed and the major organs were collected for fluorescence scanning (IVIS Lumina series 3) using 605/670 nm excitation/emission wavelength to obtain additional biodistribution data, and hematoxylin & eosin staining. Images were analyzed using Living Image Software.

## Acknowledgments

This research was co-funded by a Commonwealth Scientific and Industrial Research Organisation (CSIRO) PhD top-up scholarship. This work was performed in part at the Melbourne Centre for Nanofabrication (MCN) in the Victorian Node of the Australian National Fabrication Facility (ANFF) and Commonwealth Scientific and Industrial Research Organisation (CSIRO). The authors would also like to acknowledge Dr. Xuan Cheng from CSIRO for her assistance in TEM data acquisition. N.H.V. acknowledges funding from the Australian Research Council (in particular: IC170100016, FL220100185). A.R. acknowledges funding from the Defence Science and Technology Group (DSTG)-Australia. B.P., H.B., and W.Z. acknowledge funding from the National Natural Science Foundation of China (62288102, 62475216, 22077101)

## Table of Contents (ToC)

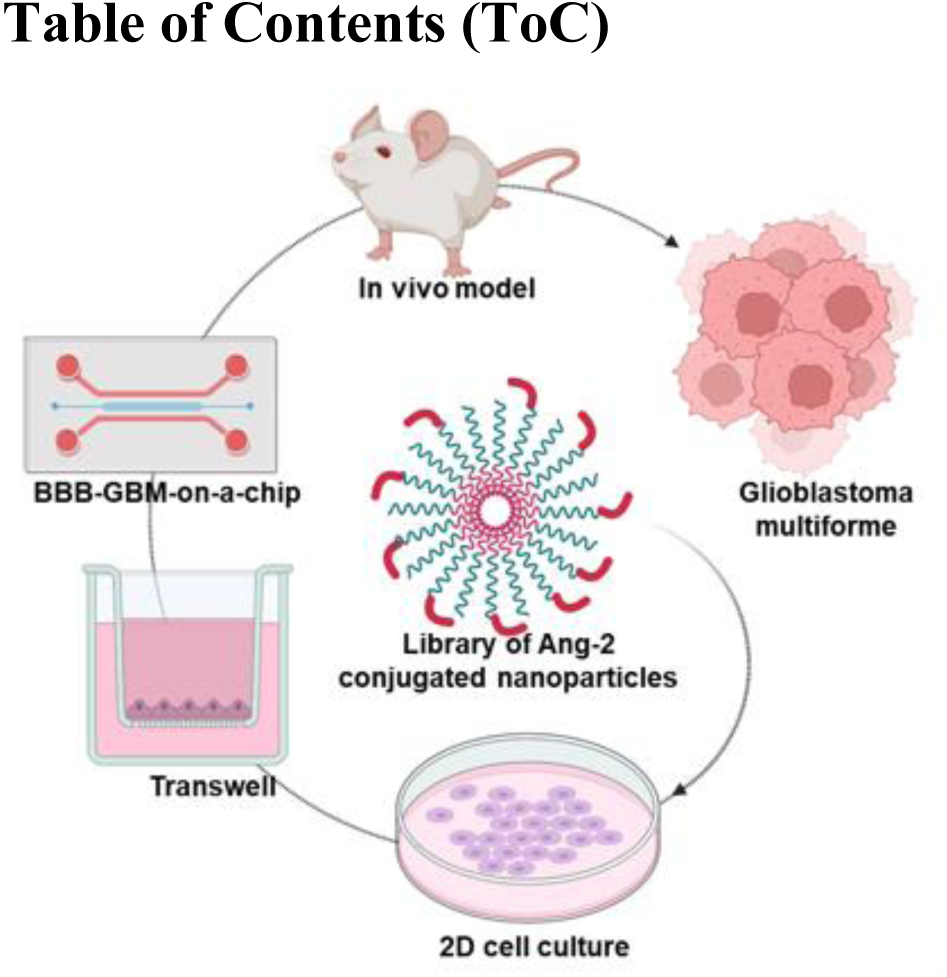

