## Supporting Information for "Optimizing Angiopep-2 Density on Polymeric Nanoparticles for Enhanced Blood-Brain Barrier Penetration and Glioblastoma Targeting: Insights from In Vitro and In Vivo Experiments"

Dr. W. Zhang, Ms. H. Li, Dr. B. Peng, Dr. H. Bai
Frontiers Science Center for Flexible Electronics, Xi'an Institute of Flexible Electronics (IFE) and Xi'an Institute of Biomedical Materials & Engineering
Northwestern Polytechnical University
127 West Youyi Road, Xi'an 710072, P. R. China

Dr. W. Zhang, Dr. A. Refaat, Ms. D. Zhu, Dr. Z. Tong, Prof. J. Nicolazzo, Dr. L. Esser, Prof. N. H. Voelcker
Drug Delivery, Disposition and Dynamics
Monash Institute of Pharmaceutical Sciences
Monash University
381 Royal Parade, Parkville, VIC 3052, Australia


Dr. W. Zhang, Dr. L. Esser
Commonwealth Scientific and Industrial Research Organisation
Research Way, Clayton, VIC 3168, Australia


Dr. A. Refaat
Pharmaceutics Department
Faculty of Pharmacy - Alexandria University
1 El-Khartoum Square, Alexandria, 21021, Egypt

Prof. N. H. Voelcker
Melbourne Centre for Nanofabrication
Victorian Node of the Australian National Fabrication Facility
151 Wellington Rd, Clayton, VIC 3168, Australia

Prof. N. H. Voelcker
Department of Materials Science & Engineering
Faculty of Engineering
Monash University
14 Alliance Ln, Clayton, VIC 3168, Australia

*** Corresponding authors: Dr. Lars Esser, and Prof. Nicolas H. Voelcker**

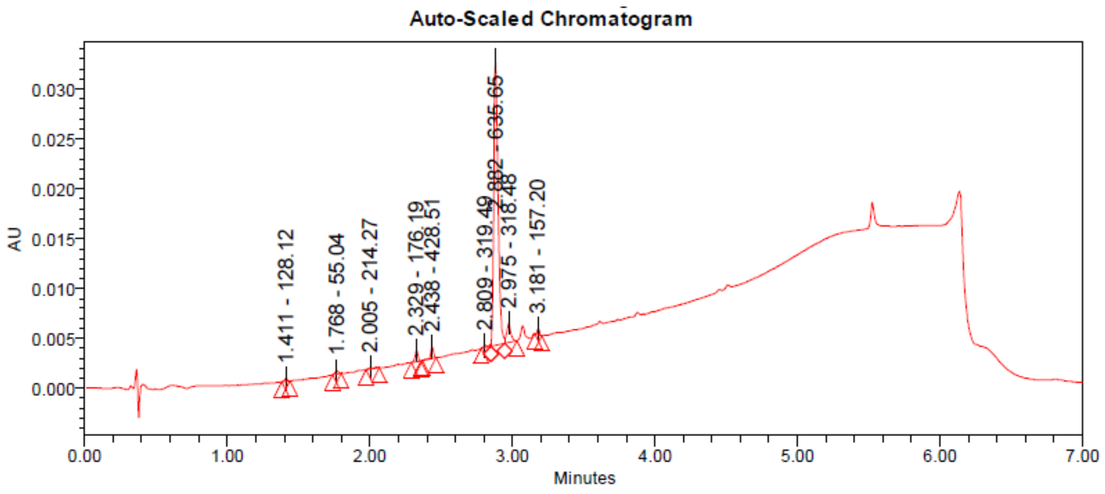


**Peak Results**

| | RT | Area | % Area | Height | Base Peak (m/z) |
|---|---|---|---|---|---|
| 1 | 1.411 | 393 | 0.56 | 293 | 128.12 |
| 2 | 1.768 | 507 | 0.72 | 330 | 55.04 |
| 3 | 2.005 | 448 | 0.63 | 160 | 214.27 |
| 4 | 2.329 | 1465 | 2.07 | 1106 | 176.19 |
| 5 | 2.438 | 1549 | 2.19 | 1178 | 428.51 |
| 6 | 2.809 | 565 | 0.80 | 255 | 319.49 |
| 7 | 2.882 | 61179 | 86.40 | 28695 | 635.65 |
| 8 | 2.975 | 4026 | 5.69 | 1972 | 318.48 |
| 9 | 3.181 | 679 | 0.96 | 602 | 157.20 |

**Figure S1**. LC-MS chromatogram of purified Cy5-acrylamide (m/z at 635.5).

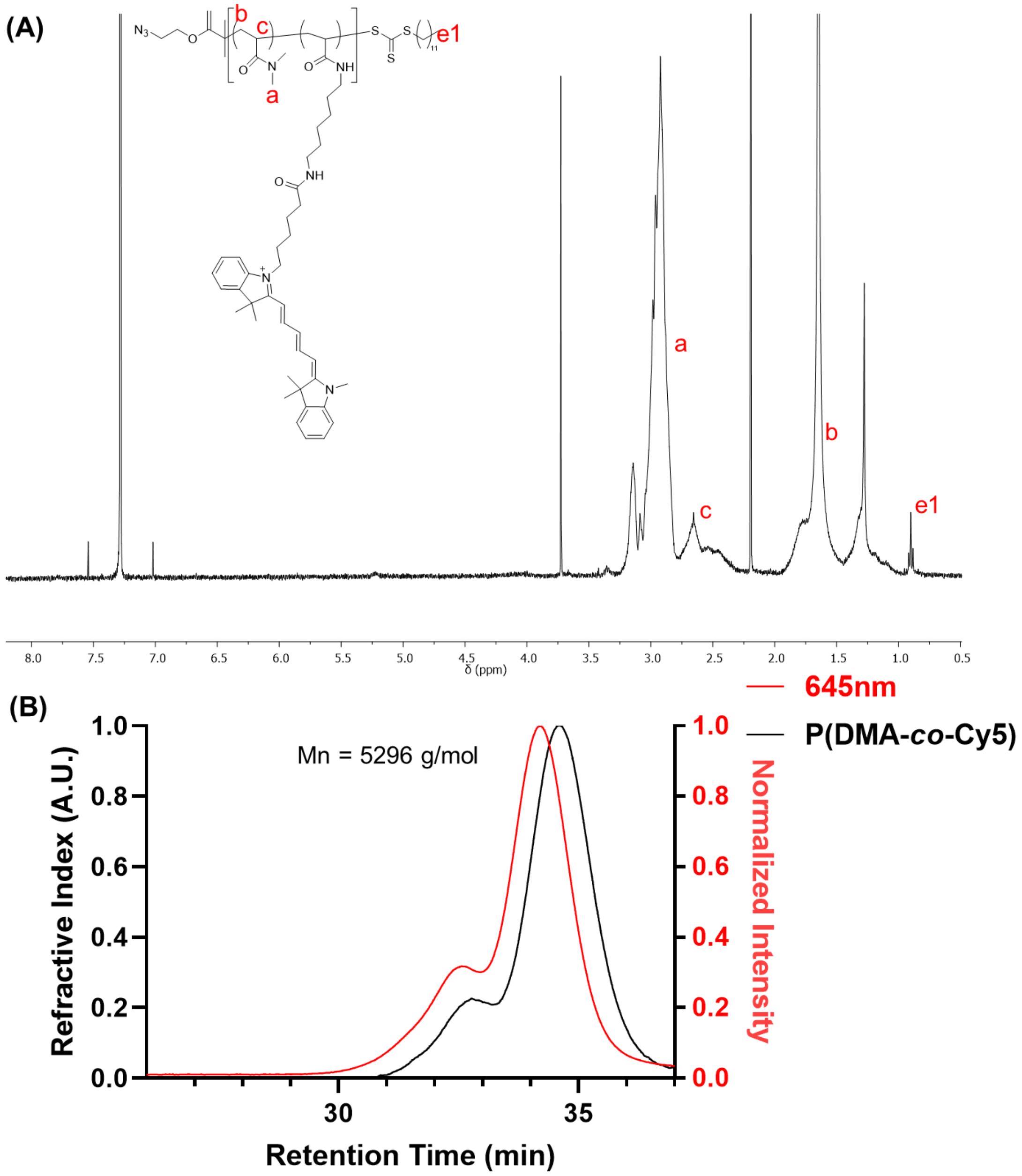


**Figure S2**. (A) $^{1}$H NMR spectrum of P(DMA-*co*-Cy5). The protons from Cy5-acrylamide are invisible due to the low feeding ratio (recorded in $CDCl_3$). (B) SEC trace with correlated fluorescence emission at 645 nm of P(DMA-*co*-Cy5).

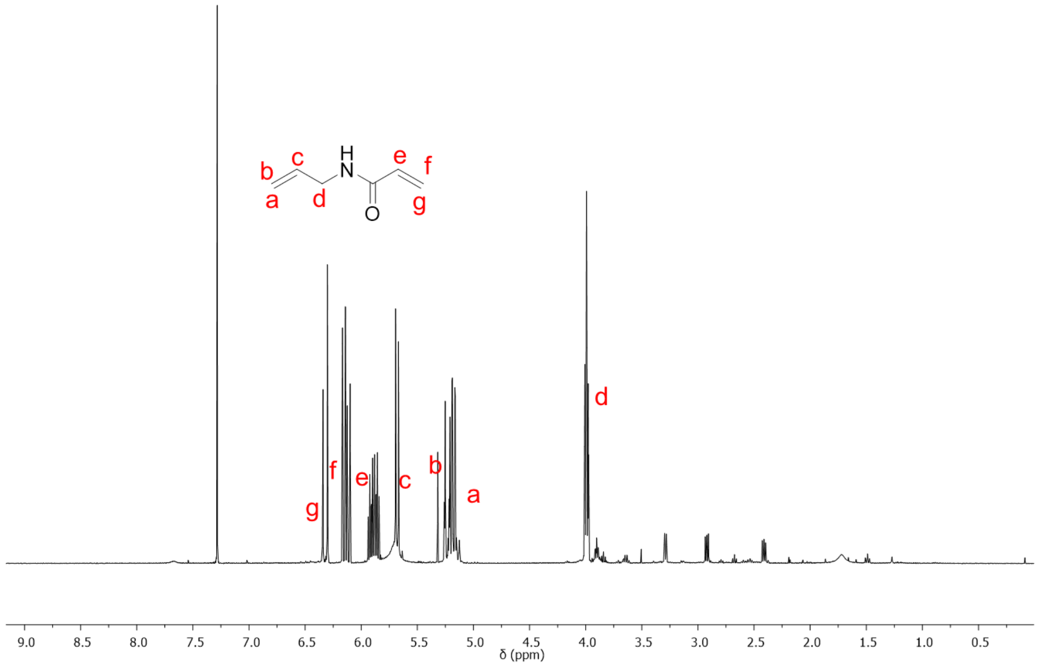


**Figure S3**. [1]H NMR spectrum of ALAM (recorded in $CDCl_3$).

| | Retention Time | Adjusted RT | Mn | Mw | MP | Mz | Mz+1 | Polydispersity | Baseline Start | Baseline End |
|---|---|---|---|---|---|---|---|---|---|---|
| 1 | 13.753 | 13.753 | 82593 | 148964 | 171901 | 214556 | 268180 | 1.803607 | 12.650 | 15.667 |

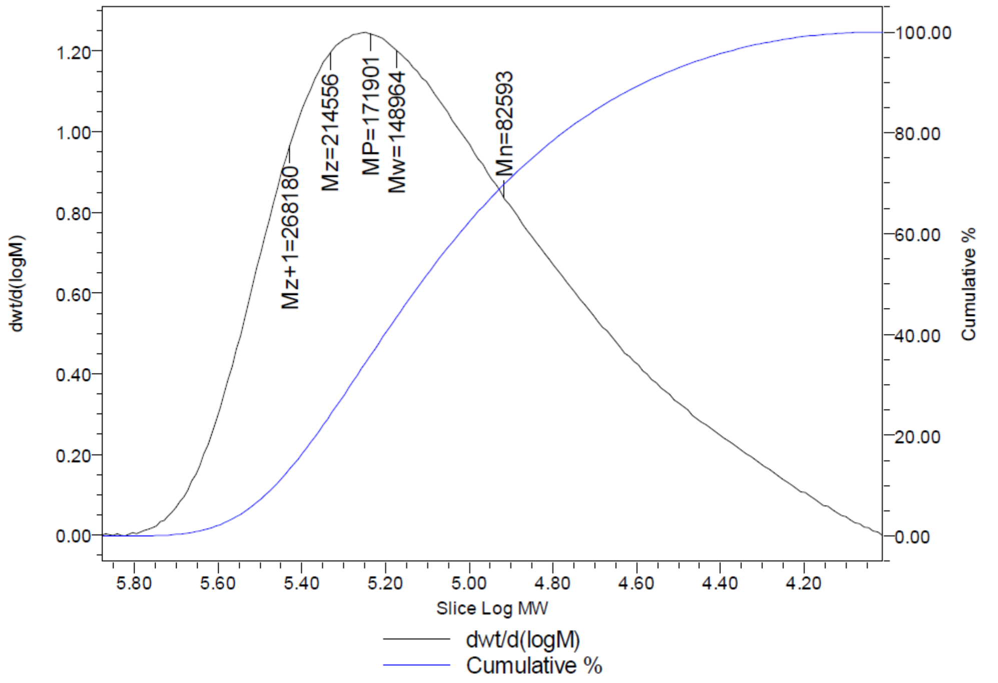


**Figure S4**. Aqueous SEC result of polymer in self-assembled nanoparticles $DAA_{100}$.

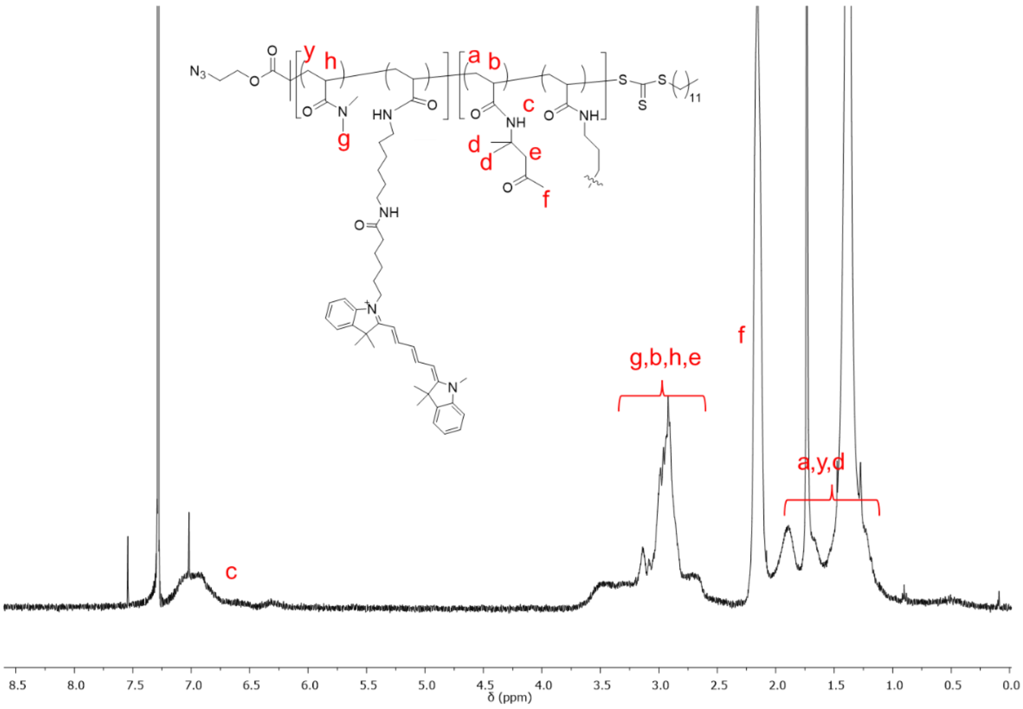


**Figure S5**. $^{1}H$ NMR spectrum of polymer in self-assembled nanoparticles $DAA_{100}$ (recorded in $CDCl_3$). The protons from Cy5-acrylamide and ALAM are invisible due to the low degree of polymerization compared to DMA and DAA.

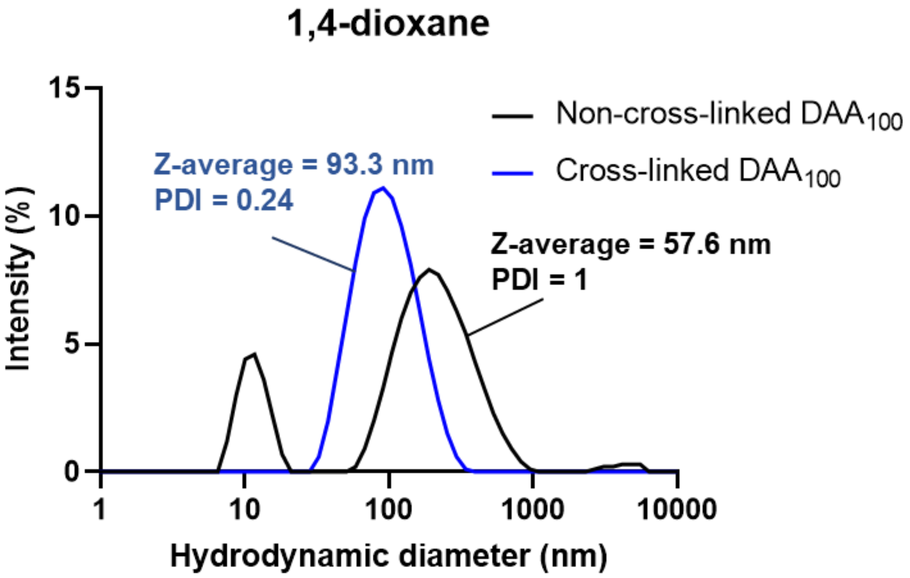


**Figure S6**. Comparison of particle hydrodynamic diameter (ζ-average) and PDI of non-cross-linked $DAA_{100}$ and cross-linked $DAA_{100}$ in the 1,4-dioxane.

| Feed molar ratio (Polymer:Ang-2) | Cy3-DBCO determined Conjugation molar ratio | Number of Ang-2 per nanoparticle | Number of Ang-2 per surface area of nanoparticle ($10^{-2}$ unit per $nm^2$) |
|---|---|---|---|
| 1:2 | 0.59 ± 0.04 | 2067.1 ± 140.1 | 9.1 ± 0.6 |
| 1:1 | 0.42 ± 0.02 | 1471.5 ± 70.1 | 6.2 ± 0.3 |
| 1:0.5 | 0.21 ± 0.02 | 745.7 ± 71.1 | 3.7 ± 0.3 |
| 1:0.1 | 0.09 ± 0.03 | 319.5 ± 106.5 | 1.5 ± 0.3 |

**Figure S7.** Estimation of Ang-2 conjugation efficiency using a complementary assay in which Cy3-DBCO was conjugated to unreacted azido-groups following Ang-2 conjugation. The polymers were first solubilised to prevent any steric hindrance that may be present when self-assembled. Values are shown as mean ± SD (n = 3).

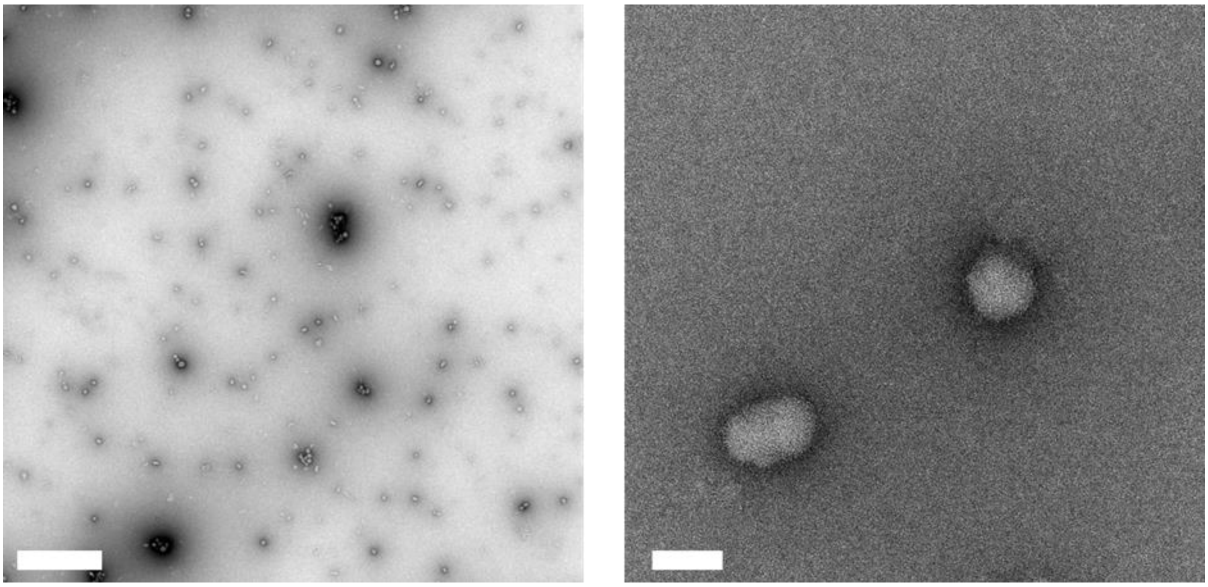

**Figure S8**. TEM images of $DAA_{100}$-Ang-2(1:2) in low (left image, scale bar = 1 μm) and high (right image, scale bar = 50 nm) magnification.

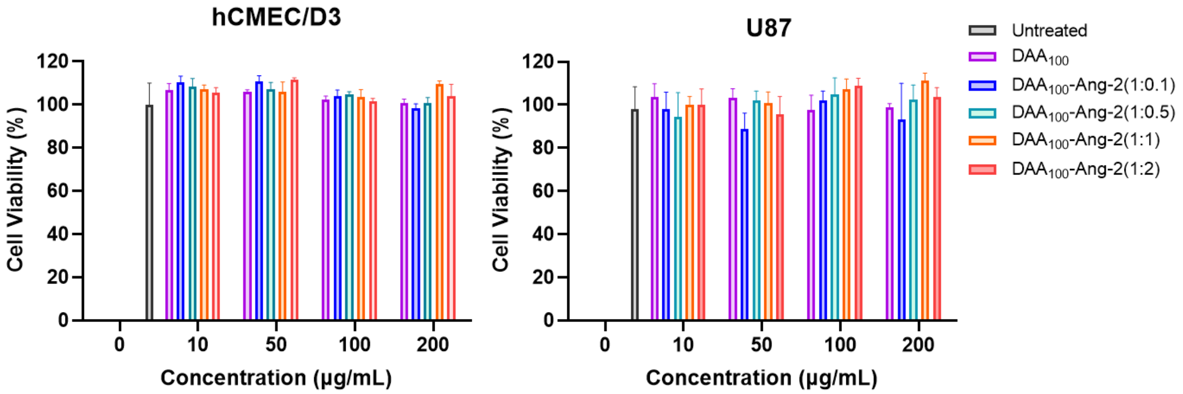


**Figure S9**. Cell viability results for hCMEC/D3 cells and U87 cells treated for 72 h with different concentrations (10, 50, 100 and 200 μg/mL) of nanoparticles with different Ang-2 conjugation ratios and evaluated using the CellTiter-Glo ATP-based luminescence cell viability assay. The values are shown as mean ± SD (n = 3).

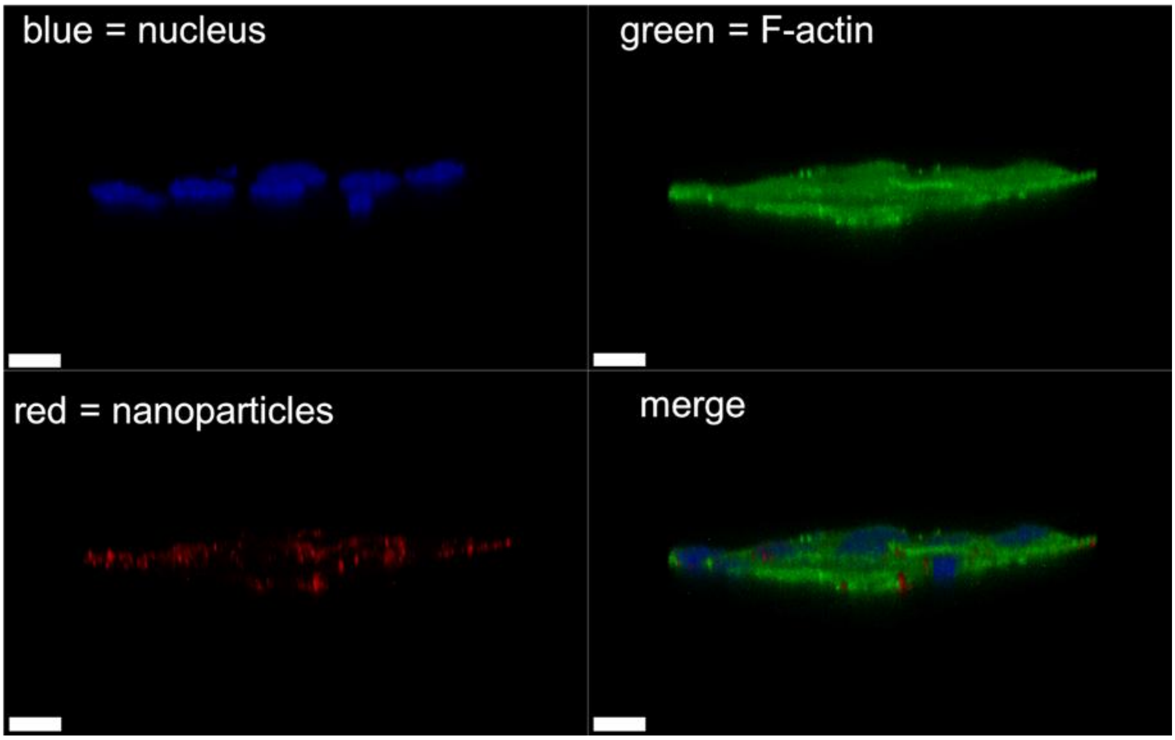


**Figure S10**. Cellular uptake of $DAA_{100}$-Ang-2(1:2) in hCMEC/D3 with z-stack scan. (blue = nucleus, green = F-actin, red = nanoparticles, scale bar = 10 μm).

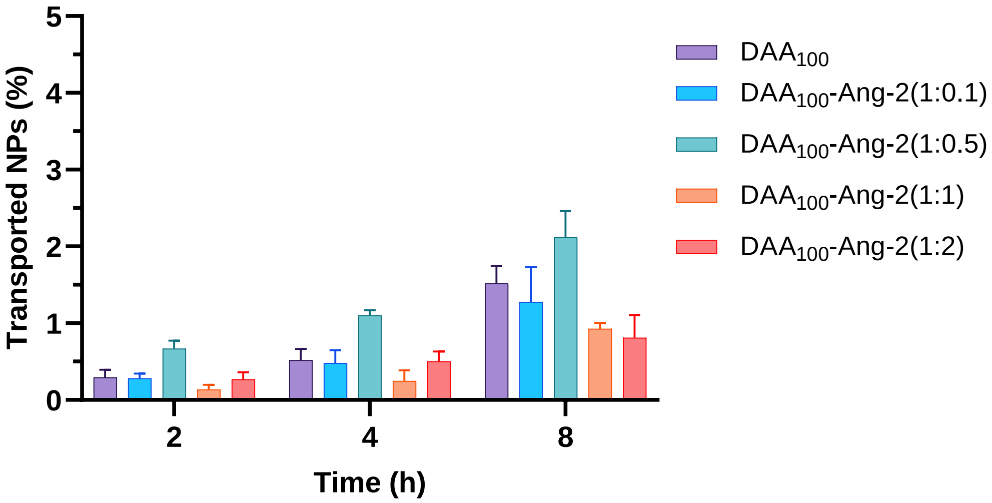


**Figure S11**. Percentage of accumulated nanoparticles *vs* time in the bottom compartment of the Transwell after applying nanoparticles in the top compartment for 2, 4 and 8 h time points. Data are shown as mean ± SD (n = 3).

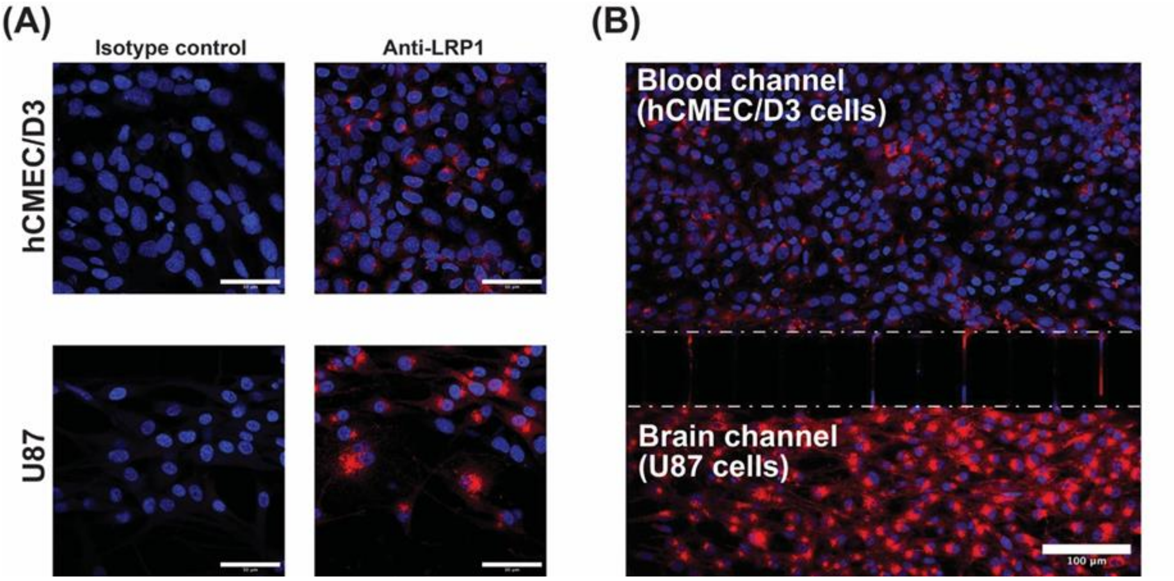


**Figure S12.** LRP1 immunostaining of hCMEC/D3 and U87 cells. Representative confocal microscopy images of (A) cells treated with Anti-LRP1 rabbit antibody, in comparison to its isotype control (scale bar = 50 µm), and (B) the BBB-GBM-on-a-chip model showing higher signals of LRP1 in the brain channel (scale bar = 100 µm). Nuclei are shown in blue while LRP1 receptors are presented in red.

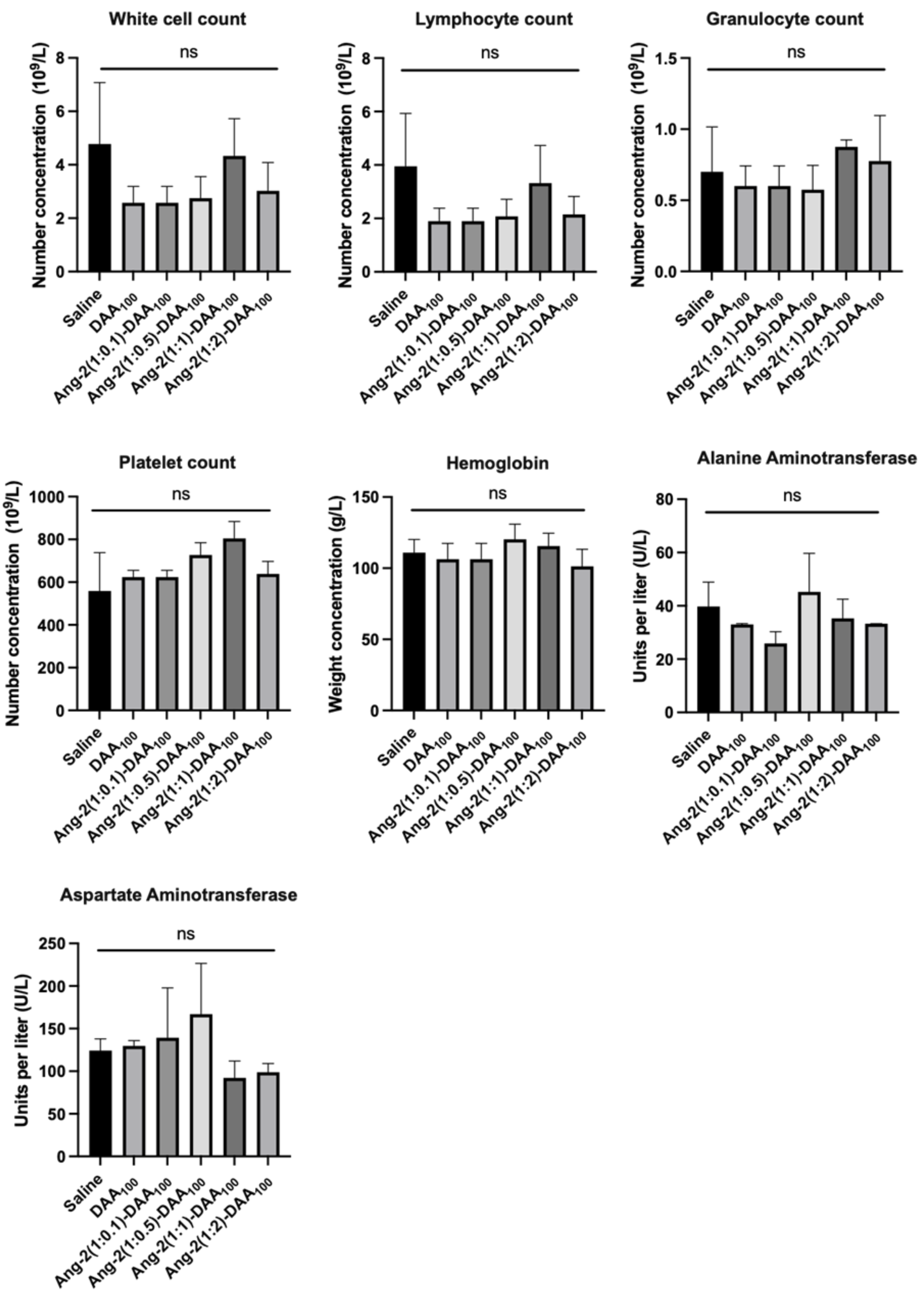


**Figure S13.** Complete blood count (CBC) and biochemical tests of alanine aminotransferase (ALT) and aspartate aminotransferase (AST) from mice following 24 h of intravenous administration of different nanoparticles at 10 mg/kg. All values are expressed as mean ± SD, n = 3, using the one-way ANOVA with Tukey's multiple comparisons test, * $p < 0.05$, ** $p < 0.01$, *** $p < 0.001$, **** $p < 0.0001$.

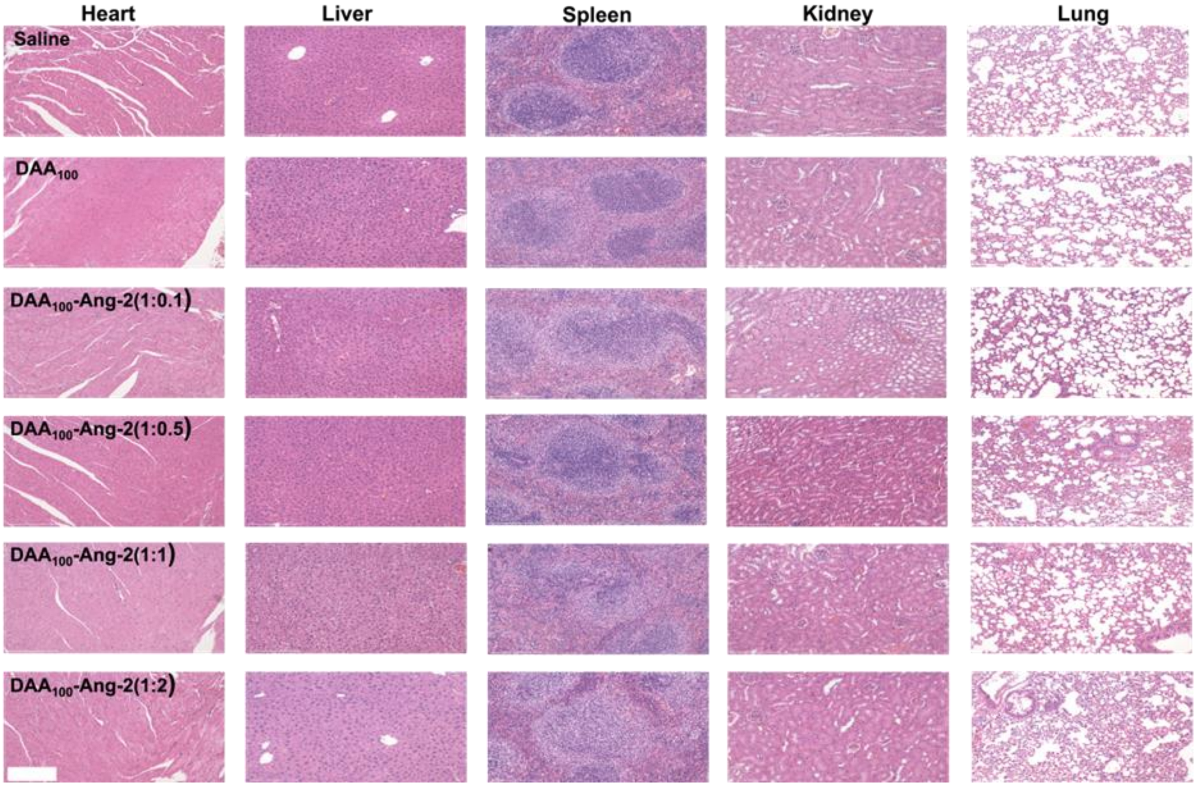


**Figure S14.** Representative H&E staining images of major organs collected from mice after 24 h of intravenous administration of nanoparticles (10 mg/kg). Scale bar, 200 μm.

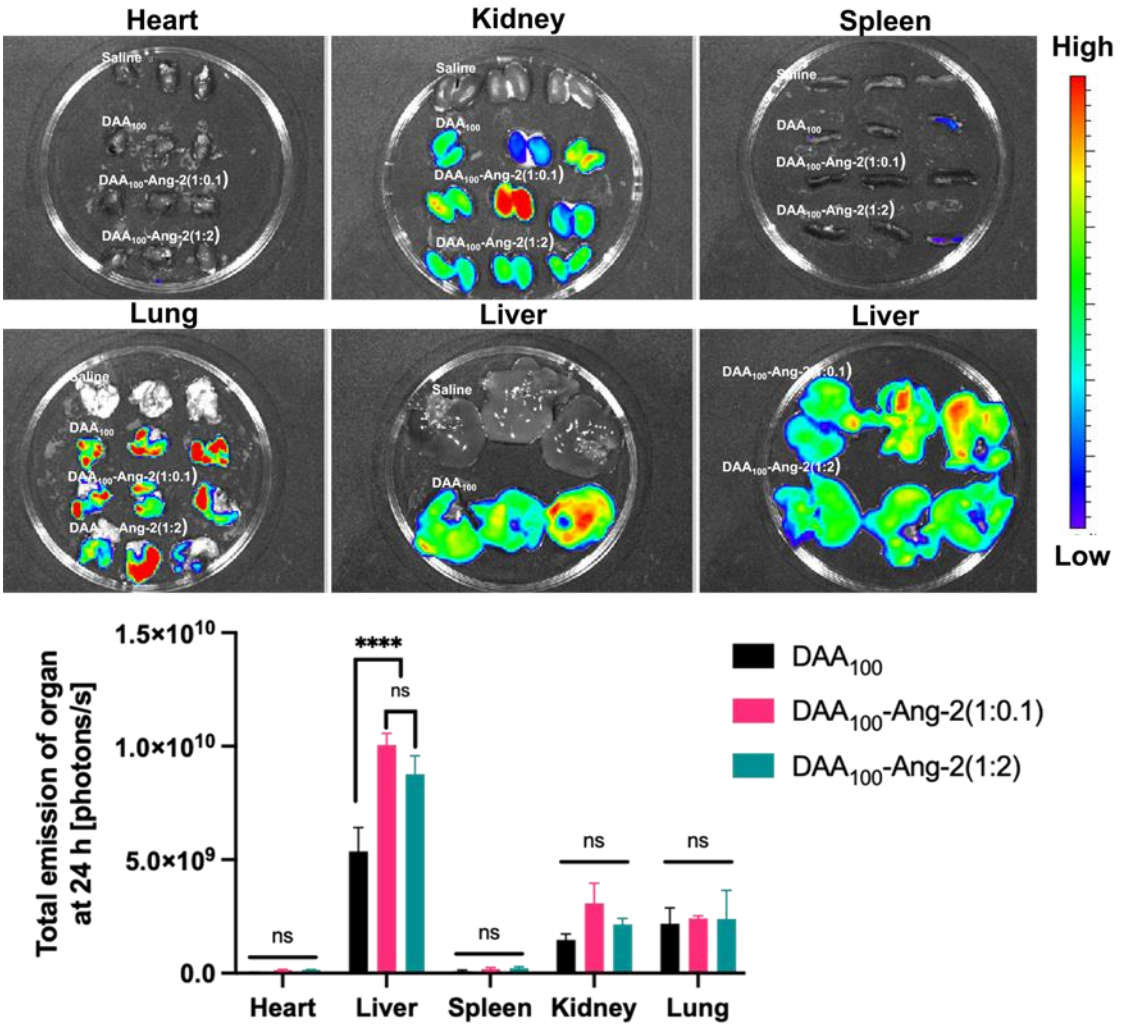


**Figure S15.** Representative *ex vivo* fluorescence scans and total emissions [photons/s] of hearts, livers, spleens, and kidneys from mice following intravenous administration at 24 h (n = 3). All values are expressed as mean ± SD, n = 3, using the two-way ANOVA with Tukey's multiple comparisons test, * $p < 0.05$, ** $p < 0.01$, *** $p < 0.001$, **** $p < 0.0001$.

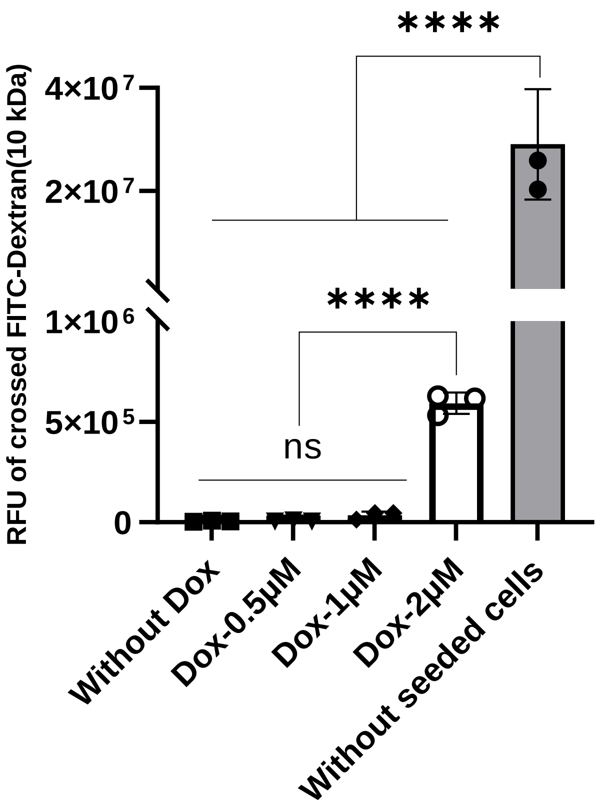


**Figure S16.** Permeation of FITC-dextran (25 µg/mL, 10 kDa) from blood to brain channels during co-administration with free Dox at concentrations of 0.5, 1, and 2 µM in an established BBB-GBM-on-a-chip. Negative control (without Dox): FITC-dextran (25 µg/mL) was flown with medium without Dox in the blood channel of the established chips. Positive control (without seeded cells): FITC-dextran (25 µg/mL) was flown with the medium in a blank chip. Corresponding relative fluorescence intensity (RFU) of FITC-dextran in brain channels in all treatments. The values are shown as mean ± SD, n= 3, using the one-way ANOVA with Tukey's multiple comparisons test, * $p < 0.05$, ** $p < 0.01$, *** $p < 0.001$, **** $p < 0.0001$.
